# Self-assembly of cellular neighborhoods converts stochastic signaling into sustained olfactory neurogenesis

**DOI:** 10.1101/2022.09.05.506659

**Authors:** Sriivatsan G. Rajan, Joseph N. Lombardo, Lynne M. Nacke, Farid Manuchehrfar, Kaelan Wong, Jocelyn Garcia, Jie Liang, Ankur Saxena

## Abstract

Olfactory neurogenesis occurs continuously throughout the lives of vertebrates, including in humans, and relies on the rapid, unceasing differentiation and integration of neurons into a complex multicellular network. The system-wide regulation of this intricate choreography is poorly understood; in particular, it is unclear how progenitor cells convert stochastic fluctuations in cell-cell signaling, over both space and time, into streamlined fate decisions. Here, we track single-cell level multicellular dynamics in the developing zebrafish olfactory epithelium, perturb signaling pathways with temporal specificity, and find that the continuous generation of neurons is driven by the spatially-restricted self-assembly of transient groups of progenitor cells, i.e. cellular neighborhoods. Stochastic modeling and validation of the underlying genetic circuit reveals that neighborhood self-assembly is driven by a tightly regulated bistable toggle switch between Notch signaling and the transcription factor Insulinoma-associated 1a that is responsive to inter-organ retinoic acid signaling. Newly differentiating neurons emerge from neighborhoods and, in response to brain-derived neurotrophic factor signaling, migrate across the olfactory epithelium to take up residence as apically-located, mature sensory neurons. After developmental olfactory neurogenesis is complete, inducing injury results in a robust expansion of neighborhoods, followed by neuroregeneration. Taken together, these findings provide new insights into how stochastic signaling networks spatially pattern and regulate a delicate balance between progenitors and their neuronal derivatives to drive sustained neurogenesis during both development and regeneration.

## INTRODUCTION

The assembly of complex neuronal structures requires the system-wide patterning and fate determination of stem cells over both space and time^1-3^. It remains unclear how cell fate decisions are influenced by a vast array of intercellular signals as stem cells migrate through a densely populated microenvironment. This intercellular communication is susceptible to a high degree of noise in cell-cell signaling and stochasticity in gene expression profiles^4-6^. *In vitro* work with 3D cell cultures and embryonic stem cell aggregates has demonstrated that seemingly homogenous progenitor cells can spontaneously self-organize into discrete domains of specialized cell types, including neurons^7-9^. However, it remains to be determined how such self-organization might direct cell fate transitions *in vivo* and how groups of progenitor cells integrate stochastic signals to direct differentiation.

The vertebrate olfactory epithelium (OE) provides a compelling model for interrogating this system-wide coordination of neurogenesis, as it harbors a high density of multiple progenitor cell types, rapid neuronal differentiation accompanied by physical expansion, and segregation into distinct apical and basal regions. The coordinated differentiation of neuronal progenitors into a diverse repertoire of olfactory sensory neurons (OSNs) is required not only during embryogenesis but well beyond, with OSNs regenerating throughout an organism’s lifetime, including in humans^10-12^. In this regard, signaling pathways that exhibit a high degree of stochasticity and cell-to-cell variation^4-6^ must somehow lead to discrete cell fate decisions not only for the initial construction of the OE’s elaborate neuroepithelial architecture but also for its continuous replenishment over a sustained period of time^13, 14^. Increasing the potential complexity of interactions, signaling inputs may converge not only from cells within the OE but from other nearby developing organs.

It remains unknown how inherent variability in the interactions between extrinsic signaling inputs and cell-intrinsic regulatory modules produces an emergent neuronal structure from a uniform progenitor field. One potential contributor is the Notch signaling pathway, which has been implicated in cell fate determination and/or differentiation in multiple tissues, including in the OE^14-18^, but with an undefined repertoire of signaling partners. Meanwhile, the transcription factor Insulinoma-associated 1a (Insm1a) has been shown to promote neurogenesis in other tissues such as the posterior lateral line, motor neurons, and photoreceptors^19-21^. In mice, Insm1 has been suggested to promote the transition of olfactory progenitors from a proliferative to differentiating state^22^, but it is unclear how and when that process is regulated and which signaling pathways might coordinate to drive neurogenesis forward.

Here, we use the developing zebrafish OE as a model to provide a unifying framework for olfactory neurogenesis that integrates signaling at multiple scales – single cells, small clusters of cells within an organ, and both intra- and inter-organ signaling. First, we quantitatively track cell signaling dynamics and employ temporally-specific perturbations to uncover a bidirectional antagonistic interaction between Notch signaling and *insm1a* that enables the coordination of continuous OSN differentiation and is dependent upon inter-organ retinoic acid (RA) signaling from the eye. To understand how several simultaneous signals translate into dynamic cell fate transitions *in vivo* in the presence of cell-to-cell variation and signaling noise, we developed a stochastic network model, whose probability landscape predicts that Notch signaling/*insm1a* mutual antagonism gives rise to divergent cell fates via a bistable toggle switch. We validate this model with quantitative imaging and analysis of relative mRNA and fluorescent reporter levels, followed by spatial mapping of basal cell distributions, revealing the self-assembly of dynamic multicellular neighborhoods. These neighborhoods consist of Notch signaling-activated ‘center’ cells surrounded by *insm1a*-expressing ‘neighbor’ cells, possess high neurogenic potential, are responsive to inter-organ RA signaling, and spawn new OSNs. Neighborhood-generated nascent neurons migrate apically towards mature OSNs via a BDNF-TrkB chemotactic pathway and integrate into the growing olfactory rosette. Finally, we demonstrate that post-development injury of the OE yields a rapid expansion of neurogenic neighborhoods, followed by OSN regeneration.

## RESULTS

### Notch signaling and *insm1a* expression are dynamic and spatially-distinct during olfactory neurogenesis

To visualize Notch signaling dynamics *in vivo* during the transition of basally-located progenitors into apically-located differentiated neurons (Fig **1A**), we performed time-lapse imaging of the developing OE in transgenic Tp1:d2GFP+ embryos (Fig **1B**, supplementary video 1), which express destabilized GFP upon Notch signaling activation^23^. At 23-31 hours post fertilization (hpf), when active olfactory neurogenesis is well underway, we individually tracked all GFP-expressing cells and found numerous basally-located cells with active Notch signaling that migrated apically within the OE (Fig **1B-B****’**, cell tracks). We observed substantial variation in cell-to-cell Notch signaling levels and found that a subset of cells exhibited robust Notch signaling activation followed by rapid downregulation and that this subset was not stereotypically conserved from embryo to embryo (Fig **1B”**). Next, we performed hybridization chain reaction (HCR)^24^ and found that mRNAs for the Notch signaling receptor *notch1a* (Fig **S1**) and its downstream effector *her4.1* are expressed predominantly in the basal part of the developing OE, where progenitors reside, at 24, 28, and 32 hpf (Fig **1C-E**). The basal localization of Tp1:d2GFP+, *notch1a*-expressing, and *her4.1*-expressing cells, as well as the variation in their expression levels from cell to cell and from embryo to embryo, cumulatively suggest that progenitor cells undergo dynamic, stochastic Notch signaling activation during this early developmental time window. By contrast, *insm1a* mRNA exhibited a spatially broader expression domain from the basal to apical regions of the OE at 24, 28, and 32 hpf (Fig **1F-H**), including in both basally-located progenitor cells and apically-located OSNs.

**Figure 1:**
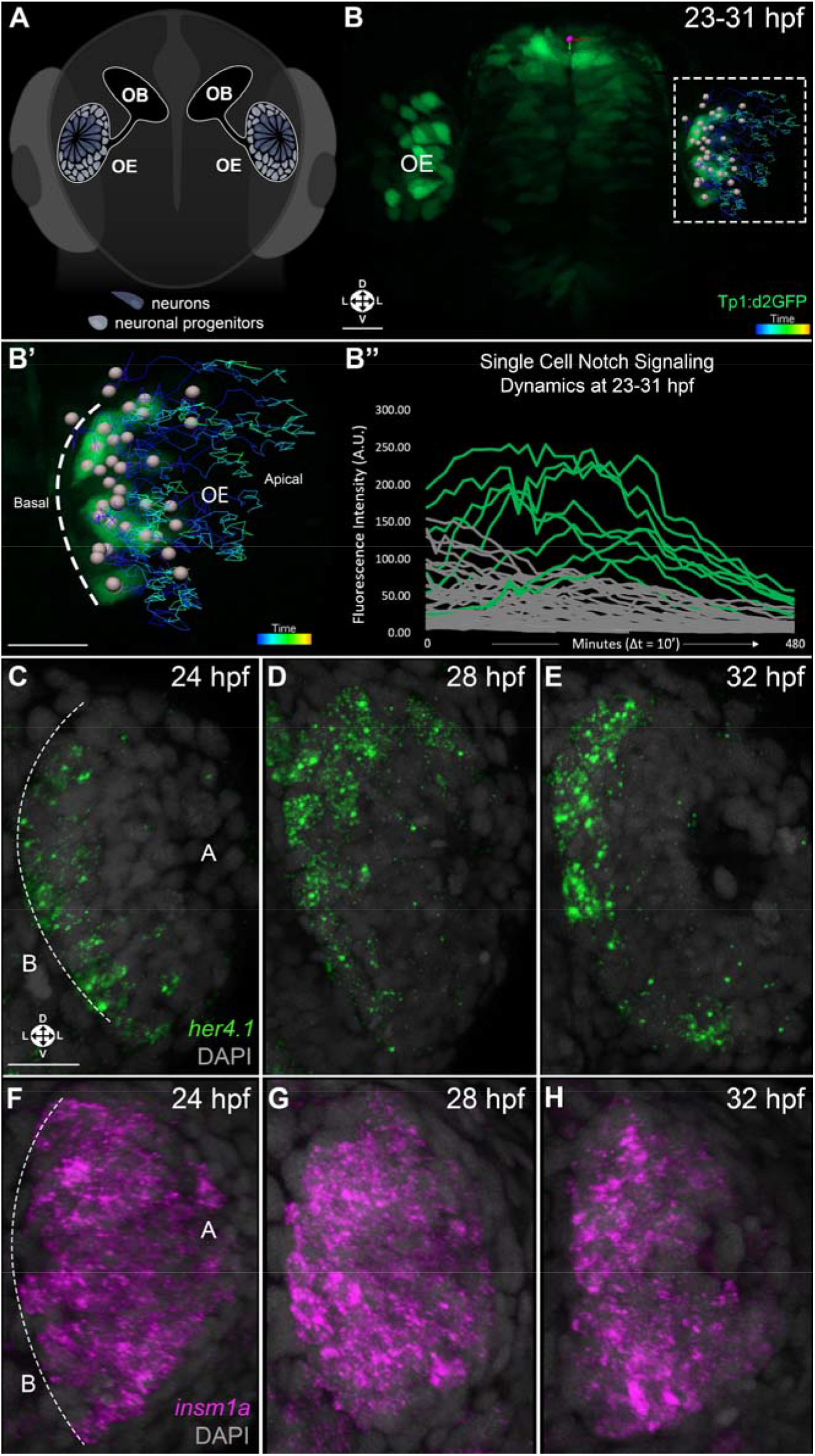
Notch signaling and *insm1a* expression are dynamic and spatially distinct during olfactory neurogenesis. (A) Schematic of zebrafish embryo (anterior view) illustrating the arrangement of progenitors and neurons in the developing olfactory epithelium (OE) (B) 3D projection anterior view of developing Tp1:d2GFP+ zebrafish embryo imaged at 23-31 hours post-fertilization (hpf), followed by tracking of d2GFP+ cells in the OE (box) (B’) Magnified image of box in B, with cell tracks demonstrating apically-directed migration over time (B’’) Quantitation of GFP signal intensity for individual migrating cells (n = 50) every 10 minutes at 23-31 hpf. Green lines indicate cells undergoing Notch signaling activation followed by downregulation. (C-H) 3D projection anterior views of whole mount *in situ* hybridization chain reaction (wmHCR) of OEs at 24 hpf (C,F), 28 hpf (D,G), and 32 hpf (E,H) showing basal expression of *her4.1* mRNA (C-E) and OE-wide expression of *insm1a* (F-H). n = 6 embryos per time point. Tp1:d2GFP: green; *her4.1* mRNA: green; *insm1a* mRNA: magenta; DAPI: grey. OE: olfactory epithelium; OB: olfactory bulb; B: basal; A: apical. Orientation arrows: L: Lateral; D: dorsal; V: ventral. Scale bars: 20 µm.

### Notch signaling and Insm1a have opposing effects on olfactory neurogenesis via mutual antagonism

The overlapping but differential expression patterns of Notch signaling components and *insm1a* suggested that they might play distinct roles in olfactory neurogenesis. To test this hypothesis, we knocked down Notch signaling in a temporally-specific manner using the heat shock-inducible transgenic zebrafish line HS:dnMAML^16^, which expresses a dominant negative form of the transcriptional coactivator Mastermind like-1 (dnMAML) and blocks downstream activation of target genes. Upon heat shock at 24 hpf, leaving earlier development unperturbed, we found that expression of the proneural gene *neurod4* was significantly increased in dnMAML+ embryos at 31 hpf in comparison to wild-type clutchmates (Fig **S2A-C**). Five hours later, at 36 hpf, we observed an increase in the number of OMP:RFP+ ciliated OSNs and Sox10:eGFP+ microvillous OSNs (Fig **2A-B’**), and we confirmed an increase in total OSN numbers with the neuron-labeling transgenic line Ngn1:nRFP (Fig **2C-E**). Finally, we assayed *insm1a* expression in Notch signaling-deficient embryos and found a significant increase in tissue-wide and per-cell *insm1a* mRNA levels prior to the increase in OSN numbers (Fig **2F-H**). Next, we perturbed Insm1a levels with translation-blocking morpholinos targeting both *insm1a* and *tp53* (to prevent non-specific cell death), as described previously^21^. Upon knocking down Insm1a, we observed a significant decrease in *neurod4* levels at 31 hpf (Fig **S2D-F**) and in the number of OSNs at 36 hpf (Fig **2I-M**). We recapitulated these results with F^0^ CRISPR/Cas9 targeting *insm1a*, paired again with the *tp53* morpholino (Fig **S3**). Insm1a/tp53-deficient embryos exhibited a significant increase in the proportion of cells with active Notch signaling at early stages of olfactory neurogenesis (Fig **2N-P**). Thus, decreased Notch signaling leads to the upregulation of *insm1a* expression, premature differentiation of neuronal progenitors, and a corresponding increase in OSN numbers, while Insm1a knockdown leads to increased Notch signaling, reduced differentiation, and a decrease in OSN numbers. Cumulatively, these results suggest that Notch signaling and Insm1a mutually antagonize each other to govern the transition from progenitor to neuron and regulate the number of OSNs during olfactory development.

**Figure 2:**
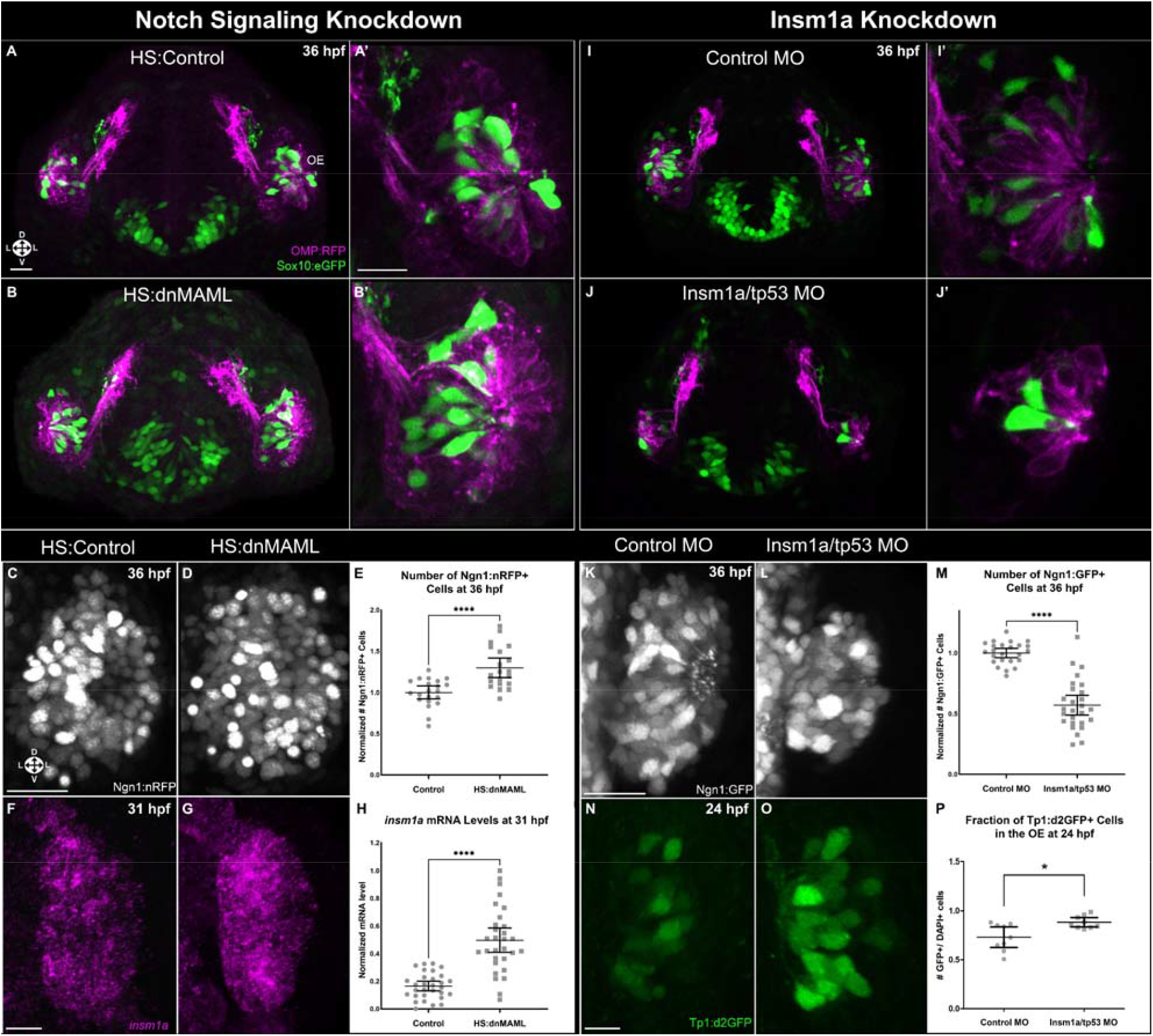
Notch signaling and Insm1a have opposing effects on neurogenesis via mutual antagonism. (A-B’) 3D projection anterior views of representative OMP:RFP+; Sox10:eGFP+ zebrafish embryos heat shocked at 24 hpf and imaged at 36 hpf without (A-A’) or with (B-B’) the heat shock-inducible transgene HS:dnMAML. Magnified optical sections (z = 20 µm) of one OE are shown in A’ and B’. (C-D) 3D projection anterior views of the OEs of Ngn1:nRFP+ embryos heat shocked at 24 hpf and imaged at 36 hpf without (C) or with (D) the heat shock-inducible transgene HS:dnMAML. (E) Normalized number of RFP+ cells at 36 hpf. Control (n = 21 embryos); HS:dnMAML (n = 20 embryos) (F-G) 3D projection anterior views of the OEs of embryos heat shocked at 24 hpf without (F) or with (G) the heat shock-inducible transgene HS:dnMAML, followed by fixation at 31 hpf and wmHCR for *insm1a* mRNA. (H) *insm1a* HCR signal intensity per cell. Number of cells: Control (n = 30 from 5 embryos); HS:dnMAML (n = 30 from 5 embryos). (I-J’) 3D projection anterior views of representative OMP:RFP+; Sox10:eGFP+ zebrafish embryos injected with control morpholino (I-I’) or Insm1a/tp53 morpholino (J-J’) and imaged at 36 hpf. (K-L) 3D projection anterior views of the OEs of Ngn1:GFP+ embryos post-injection with control morpholino (K) or Insm1a/tp53 morpholino (L) and imaged at 36 hpf. (M) Normalized number of GFP+ cells per embryo at 36 hpf. Control morpholino (n = 24 embryos); Insm1a/tp53 morpholino (n = 27 embryos). (N-O) 3D projection anterior views of the OEs of Tp1:d2GFP+ embryos post-injection with control morpholino (N) or Insm1a/tp53 morpholino (O) and imaged at 24 hpf. (P) GFP+:DAPI+ ratios in the OE. Control morpholino (n = 9 embryos); Insm1a/tp53 morpholino (n = 9 embryos). Horizontal bars: mean and 95% confidence intervals. Statistical significance: *p < 0.05, **p < 0.01, ***p < 0.001, ****p < 0.0001. Sox10:eGFP: green; OMP:RFP: magenta; Ngn1:nRFP: white; Ngn1:GFP: white; Tp1:d2GFP: green; *insm1a* mRNA: magenta. OE: olfactory epithelium. Orientation arrows: L: Lateral; D: dorsal; V: ventral. Scale bars: 20 µm.

### Inter-organ retinoic acid signaling modulates Notch signaling/Insm1a-mediated neurogenesis

We have a limited understanding of how large-scale RA gradients impact cell-cell interactions to influence tissue-wide outcomes, including in the developing OE. Previous work using mouse embryos and organ explants demonstrated that mesenchymal RA signaling is critical for induction of the olfactory placode^25^, but others have found this early requirement to be dispensable, while instead suggesting a later inhibitory role for RA in the induction of neuronal progenitors^13^. It thus remains unclear how RA might affect olfactory neurogenesis *in vivo* and, in particular, how these signaling gradient-mediated effects might vary at different times and locations, including in the rapidly expanding basal OE where olfactory progenitors reside.

In zebrafish, RA is synthesized by the retinaldehyde dehydrogenases aldh1a2 and aldh1a3 and metabolized by the catabolic enzyme cyp26a1, which, in turn, is induced by RA^26, 27^. We performed HCR for *aldh1a2, aldh1a3*, and *cyp26a1* at 31 hpf and found that *aldh1a2* and *aldh1a3* are expressed primarily in the developing eye but not in the OE (Fig **3A**). By contrast, *cyp26a1* expression is localized to just a few medially-located cells in the nearby OE at 31 hpf and at additional time points (Fig **3A-A’**, **S4**), suggesting that an RA ‘sink’ is present in between basally-located progenitor cells and apically-located OSNs. We blocked RA production at 24-31 hpf with the chemical inhibitor DEAB and found that both the number of *cyp26a1*+ cells and per cell mRNA levels of *cyp26a1* are significantly decreased in the OEs of DEAB-treated embryos (Fig **3A-D**, supplementary video 2), consistent with a reduced need for an RA sink upon reduced RA signaling. We also found an increase in *her4.1* levels, a decrease in *insm1a* levels, and a decrease in *neurod4* levels on a per cell basis in the basal region where neuronal progenitors reside (Fig **3E-G**), suggesting that RA signaling normally promotes neuronal commitment by tipping the Notch signaling/*insm1a* balance towards higher *insm1a* expression. To determine if RA synthesis in the eye and its diffusion to the nearby OE was sufficient to alter this balance, we placed RA-soaked beads into the developing eye during DEAB incubation and found that this treatment was able to rescue the effects of RA synthesis inhibition (Fig **S5**). Together, these data demonstrate that inter-organ RA signaling from the eye to the nose promotes the neuronal differentiation of basal olfactory progenitors by altering tightly regulated Notch signaling/Insm1a mutual antagonism.

**Figure 3:**
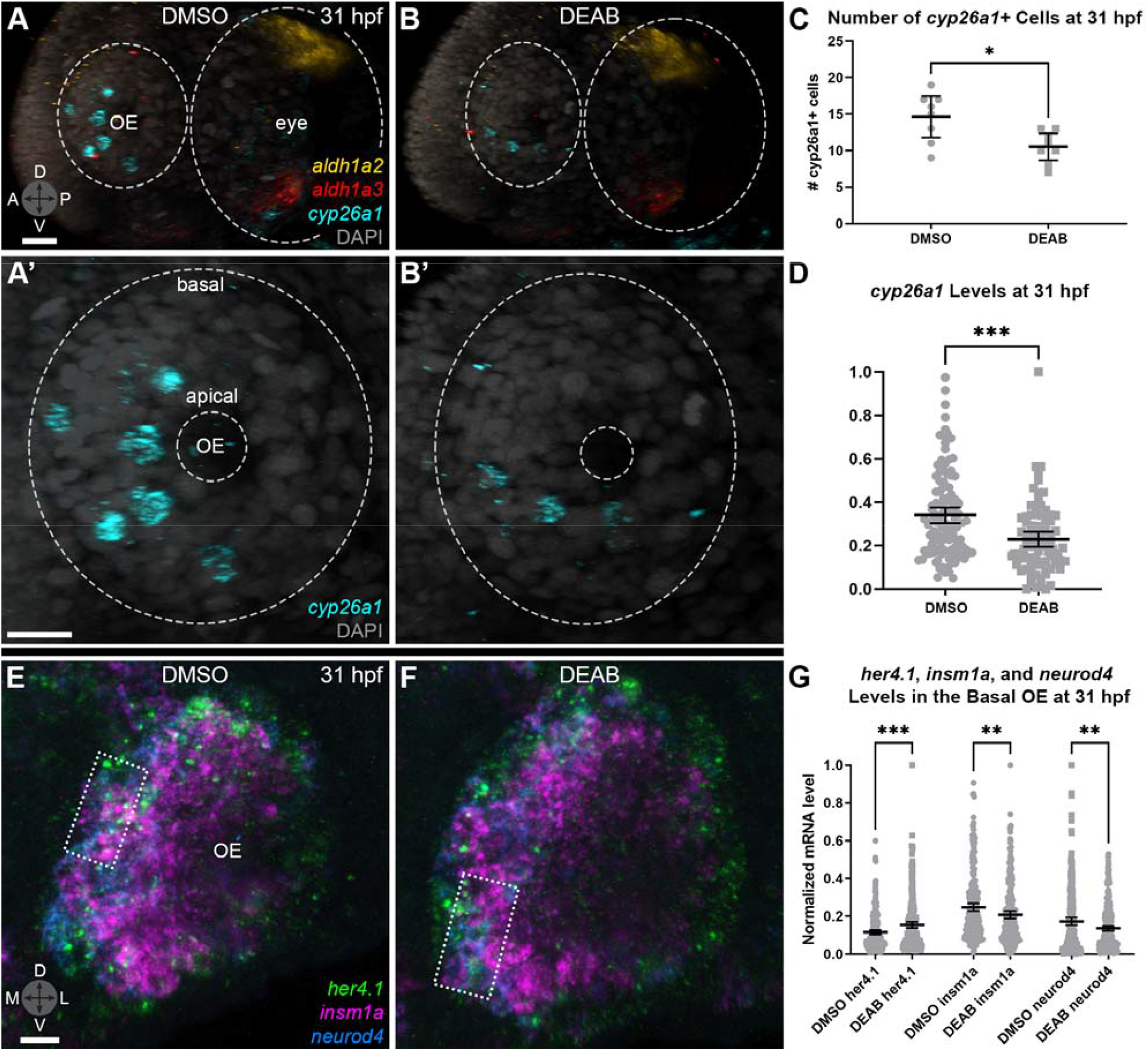
Inter-organ retinoic acid (RA) signaling alters Notch signaling/Insm1a balance. (A-B’) 3D projection lateral views of representative embryos treated with DMSO (negative control; A) or 10 µM DEAB/DMSO (RA synthesis inhibitor; B) from 24 to 31 hpf, followed by fixation at 31 hpf and wmHCR targeting *aldh1a2, aldh1a3*, and *cyp26a1* mRNA. Dashed lines outline the OE and eye. (A’,B’) Magnified views of OEs from (A,B). (C) Number of *cyp26a1*+ cells per embryo and (D) *cyp26a1* levels per cell at 31 hpf. Number of cells: DMSO (n = 117 from 8 embryos); DEAB (n = 84 from 8 embryos). (E,F) 3D projection anterior views of representative embryos treated with DMSO (E) or 10 µM DEAB/DMSO (F) from 24 to 31 hpf, followed by fixation at 31 hpf and wmHCR targeting *her4.1, insm1a*, and *neurod4* mRNA. (G) *her4.1, insm1a*, and *neurod4* mRNA levels per cell in the basal OE at 31 hpf. Number of cells: DMSO (n = 272 from 3 embryos); DEAB (n = 276 from 3 embryos). Horizontal bars: mean and 95% confidence intervals. Statistical significance: *p < 0.05, **p < 0.01, ***p < 0.001. *aldh1a2*: yellow; *aldh1a3*: red; *cyp26a1*: cyan; DAPI: grey; *her4.1*: green; *insm1a*: magenta; *neurod4*: blue. Orientation arrows: D, dorsal; V, ventral; A, anterior; P, posterior; M, medial; L, lateral. Scale bars: (A,A’) 30 µm; (E) 10 µm.

### A Notch signaling/Insm1a bistable toggle switch yields self-assembled cellular neighborhoods

Our data demonstrated that the expression of *her4.1* and *insm1a* is dependent on highly dynamic Notch signaling and RA signaling, but the artificially severe gain/loss-of-function changes introduced by our experimental perturbations are far greater than the natural fluctuations that exist in cells, as shown in both our experiments (Fig **1B****’’**, **3A-A’**) and previously in other experimental systems^28, 29^. To understand how naturally occurring variability in these signaling inputs yields starkly different outputs of either progenitor cells or differentiating neurons, we built a principled model that accounts for cell-to-cell variation in gene expression and the stochastic effects found in signal transduction. This stochastic network model was constructed using our experimental data, and its behavior was specified by its time-evolving and steady state probability landscapes, which were computed using the method of Accurate Chemical Master Equation (ACME)^30-36^. We modeled Notch and RA signaling-dependent interactions between *her4.1 and insm1a* as a mutually antagonistic toggle switch system with multiple inputs (Fig **4A**, **S6A**) and computed the probability landscape of the system, identifying a multistability regime at the steady state that overlaps substantially with the observed distribution of cells at 36 hpf (Fig **4B-C**). To determine if this stochastic network model with a multistable configuration could predict the probability landscapes at earlier time points, we generated a dynamic simulation of time series and found shifting probability landscapes that correlated substantially with our observed basal cell distributions at 24, 28, and 32 hpf (Fig **4D**, supplementary video 3). Additionally, we found that regions of high probability peaks indicating Her4.1^ON^Insm1a^OFF^ and Her4.1^OFF^Insm1a^ON^ network states (Fig **4B**) correspond to a bistable distribution of cells exhibiting *her4.1*^HIGH^*insm1a*^LOW^ and *her4.1*^LOW^*insm1a*^HIGH^ expression profiles, respectively, from 24 hpf to 36 hpf (Fig **4C-D**), suggesting the operation of a bistable toggle switch system. Going back to our *in vivo* imaging data, we identified these cells and discovered a recurring spatial pattern of *her4.1*^HIGH^*insm1a*^LOW^ ‘center’ cells in contact with ‘neighboring’ *her4.1*^LOW^*insm1a*^HIGH^ cells, together forming well-defined cellular neighborhoods in the basal region of the OE (Fig **4E**, **S6B**). We confirmed this recurring self-assembly of neighborhoods with the Notch signaling reporter Tp1:d2GFP and an *insm1a* expression reporter insm1a:mCherry (Fig **S6C**). Strikingly, the pattern of center and neighbor cells was conserved at multiple time points throughout peak periods of olfactory neuronal differentiation (Fig **4F**), suggesting that neighborhoods are critical for sustaining olfactory neurogenesis. Thus, *in silico* modeling and its cross-validation *in vivo* point to a multistable system across all basal cells. This multistability narrows down, within a subset of cells, to function as a bistable Notch signaling/Insm1a toggle switch that drives the self-assembly of cellular neighborhoods.

**Figure 4:**
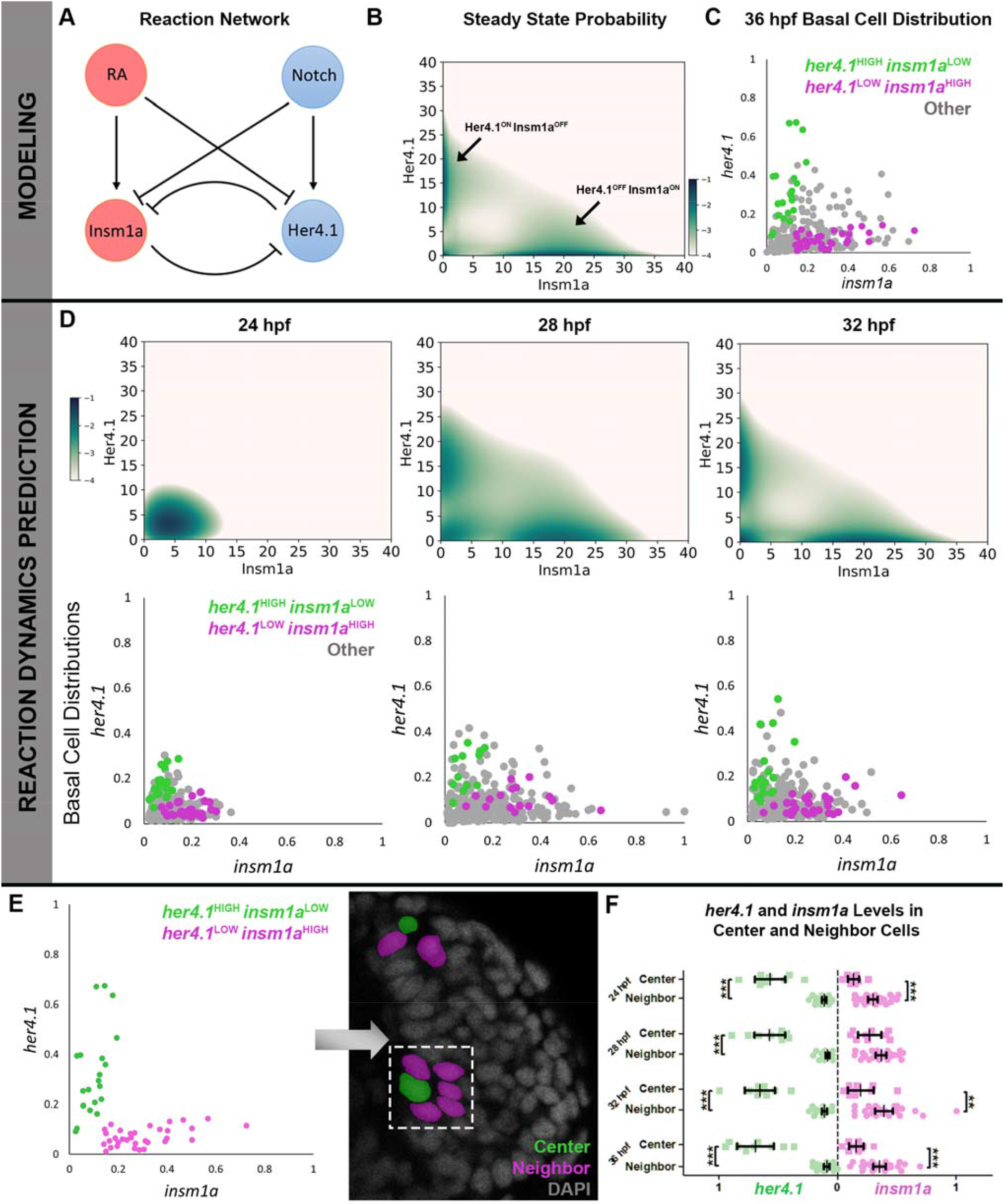
A Notch signaling/Insm1a bistable toggle switch yields self-assembled cellular neighborhoods. (A) Reaction network model and (B) steady state probability landscape plotted on a natural log scale with regions of high probability in green and regions with probability close to zero in white. Y and X axes indicate protein levels for Her4.1 and Insm1a, respectively, on a scale of 0 (minimum) to 40 (maximum). Arrows indicate two peaks for Her4.1^ON^Insm1a^OFF^ and Her4.1^OFF^Insm1a^ON^ system states. (C) Distribution of all basal cells in the OE at 36 hpf based on relative mRNA levels of *her4.1* and *insm1a* normalized to *b-actin* mRNA levels, measured via HCR. Expression profiles of subsets of cells are indicated as: *her4.1*^HIGH^*insm1a*^LOW^ (green); *her4.1*^LOW^*insm1a*^HIGH^ (magenta); other cells (grey) (D) Prediction of reaction network dynamics via Accurate Chemical Master Equation (ACME) computation. Predicted sequential probability landscapes (top row) are similar to actual basal cell distributions (bottom row) at 24, 28, and 32 hpf. Number of cells: 24 hpf (n = 302); 28 hpf (n = 273); 32 hpf (n = 296); 36 hpf (n = 301) from 3 embryos per time point. (E) Cells with reciprocal *her4.1*^HIGH^*insm1a*^LOW^ (green) and *her4.1*^LOW^*insm1a*^HIGH^ (magenta) expression exhibit a bistable configuration and are spatially arranged in a neighborhood pattern. *her4.1*^HIGH^*insm1a*^LOW^: green; *her4.1*^LOW^*insm1a*^HIGH^: magenta. For all graphs of basal cell distributions (C-E), Y and X axes indicate normalized mRNA levels for *her4.1* and *insm1a*, respectively, on a scale of 0 (minimum) to 1 (maximum). (F) Relative *her4.1* and *insm1a* mRNA levels per cell based on HCR fluorescence intensity in center and neighbor cells at 24, 28, 32, and 36 hpf. Number of cells: 24 hpf (n = 39); 28 hpf (n = 38); 32 hpf (n = 36); 36 hpf (n = 37) from 3 embryos per time point. Vertical bars: mean and 95% confidence intervals. Statistical significance: ns (not significant) p > 0.05, **p < 0.01, ***p < 0.001.

### Cellular neighborhoods continuously produce OSNs

To test the prediction that center and neighbor cells have different cellular fates, we assayed for levels of the neural stem cell marker *ascl1a* and the neuronal progenitor marker *neurod4*. At 31 hpf, center cells expressed high levels of *ascl1a* in comparison to neighbor cells (Fig **5A-B**, arrowhead and asterisk), while both center and neighbor cells expressed similar levels of *neurod4* (Fig **5C-D**, arrowhead and asterisk). Meanwhile, cells located outside of neighborhoods expressed lower levels of *ascl1a* and *neurod4* in comparison to center cells and neighborhood cells, respectively (Fig **5A-D**), suggesting that cells in neighborhoods have a markedly higher neurogenic potential than those located elsewhere (Fig **5E**). To directly determine whether neighborhoods give rise to OSNs, we performed time-lapse imaging of Tp1:d2GFP; Ngn1:nRFP transgenic embryos (Fig **S7A**) and observed a downregulation in Notch signaling as well as a concomitant increase in Ngn1:nRFP over time in Tp1:d2GFP+ cells (Fig **S7B**). We similarly imaged Tp1:d2GFP; insm1a:mCherry embryos and identified center cells transitioning into neighbor cells prior to neuronal differentiation (Fig **S8A-C**, arrowheads). Finally, time-lapse imaging of Ngn1:GFP; insm1a:mCherry embryos demonstrated insm1a:mCherry+ neighbor cells’ neuronal differentiation (Fig **S8D-D****’**). Thus, neighborhood-based cells continuously emerge and differentiate into OSNs, with a center to neighbor to neuron series of transitions.

**Figure 5:**
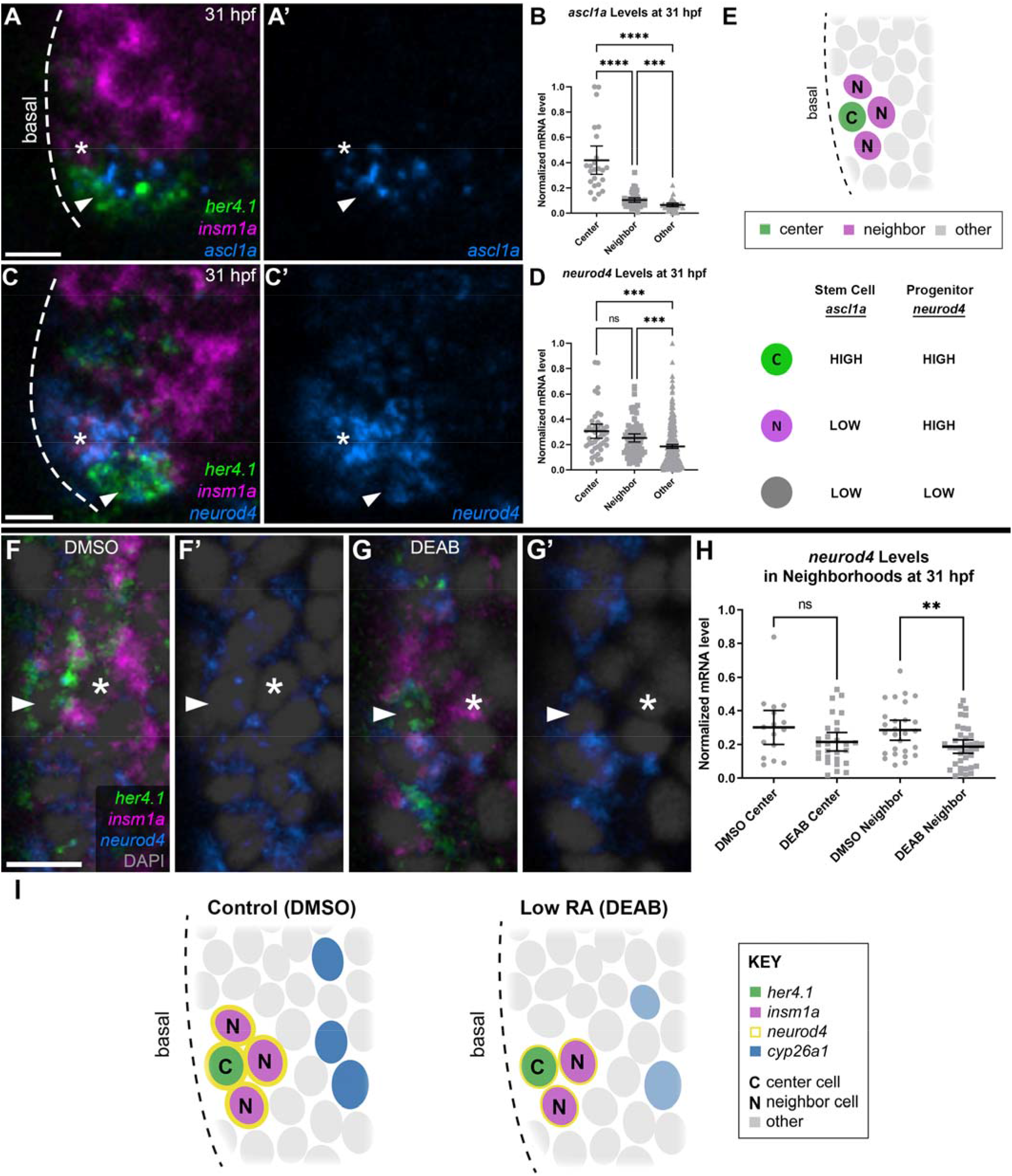
Cellular neighborhoods give rise to OSNs. (A-A’) Optical sections (z = 1.25 µm) of representative embryos fixed at 31 hpf, followed by wmHCR with probes targeting *her4.1, insm1a*, and *ascl1a* mRNA. (B) *ascl1a* mRNA levels per cell in the basal OE in center, neighbor, or other cells at 31 hpf. Number of cells: (n = 107 from 6 embryos). (C-C’) Optical sections (z = 1.25 µm) of representative embryos fixed at 31 hpf, followed by wmHCR with probes targeting *her4.1, insm1a*, and *neurod4* mRNA. (D) *neurod4* mRNA levels per cell in the basal OE in center, neighbor, or other cells at 31 hpf. Number of cells: (n = 542 from 6 embryos). (E) Schematic of neighborhood composition and stem/progenitor cell marker expression. (F-G’) Optical sections (z = 1.25 µm) of representative embryos treated with DMSO (F-F’) or 10 µM DEAB/DMSO (G-G’) at 24-31 hpf and fixed at 31 hpf, followed by wmHCR targeting *her4.1, insm1a*, and *neurod4* mRNA. Examples of center cells (arrowheads) and neighbor cells (asterisks) are indicated. (H) *neurod4* mRNA levels per cell in center and neighbor cells at 31 hpf. Number of cells: DMSO (n = 272 from 3 embryos); DEAB (n = 276 from 3 embryos). (I) Schematic of the effects of RA on neighborhoods. Horizontal bars: mean and 95% confidence intervals. Statistical significance: ns (not significant) p > 0.05, **p < 0.01, ***p < 0.001, ****p < 0.0001. *her4.1*: green; *insm1a*: magenta; DAPI: grey; Tp1:d2GFP: green; insm1a:mCherry: magenta; *ascl1a*: blue; *neurod4*: blue. Scale bars: 10 µm.

We next assayed the effect of reduced RA synthesis on neighborhoods and found a significant reduction in DEAB-treated neighbor cells’ levels of *neurod4* (Fig **5F-H**, arrowheads, asterisks) and a decrease in OSN numbers at 31 hpf that, after ceasing DEAB treatment, quickly recovered by 36 hpf (Fig **S9**). These results suggest that inter-organ RA signaling tightly regulates the composition and output of neurogenic neighborhoods, and medially-located *cyp26a1*-expressing cells (Fig **3A-A**’, **S4**, **5I**) likely divide the developing OE into zones of high RA (basal) and low RA (apical), governing the availability of RA and its effect on neighborhoods.

### Neuronal BDNF facilitates basal to apical migration of progenitors via TrkB signaling

Having observed neighborhood-based cells emerge, migrate apically, and differentiate into OSNs (Fig **1B’**, **S7A**), we wondered what was directing this continuous migration across the growing OE. Cultured human olfactory progenitors have been found to migrate in response to BDNF *in vitro*^37^, and we had previously observed highly localized *bdnf* mRNA expression in the OE at earlier time points^38^. Performing HCR at time points of peak neurogenesis, we found *bdnf* expressed specifically in apically-located differentiated neurons at 32 hpf (Fig **6A-A****’**) and its receptor, TrkB orthologue *ntrk2a*, expressed in basally-located neighborhoods (Fig **6B-B****’**). Hypothesizing that newly differentiating neurons exit basally-located neighborhoods and migrate along an increasing gradient of OSN-secreted BDNF, we treated Tp1:d2GFP+ embryos with DMSO (Fig **6C-C****’**) or BDNF-TrkB signaling inhibitor ANA-12 (Fig **6D-D****’**) starting at 22 hpf, performed time-lapse imaging at 26-36 hpf, and tracked Tp1:d2GFP+ cells’ movements in the OE (Fig **6E-F**). BDNF-TrkB signaling inhibition did not affect cellular motility (Fig **6E**) but did result in a significant reduction of directed migration along the radial axis towards the apical region (Fig **6C-D****’, F**). These data suggest that production of BDNF by existing OSNs (Fig **6A-A****’)** guides the directed basal-to-apical movement of TrkB-expressing differentiating neurons (Fig **6B-B****’**, inset) and facilitates their integration into the expanding neuronal rosette.

**Figure 6:**
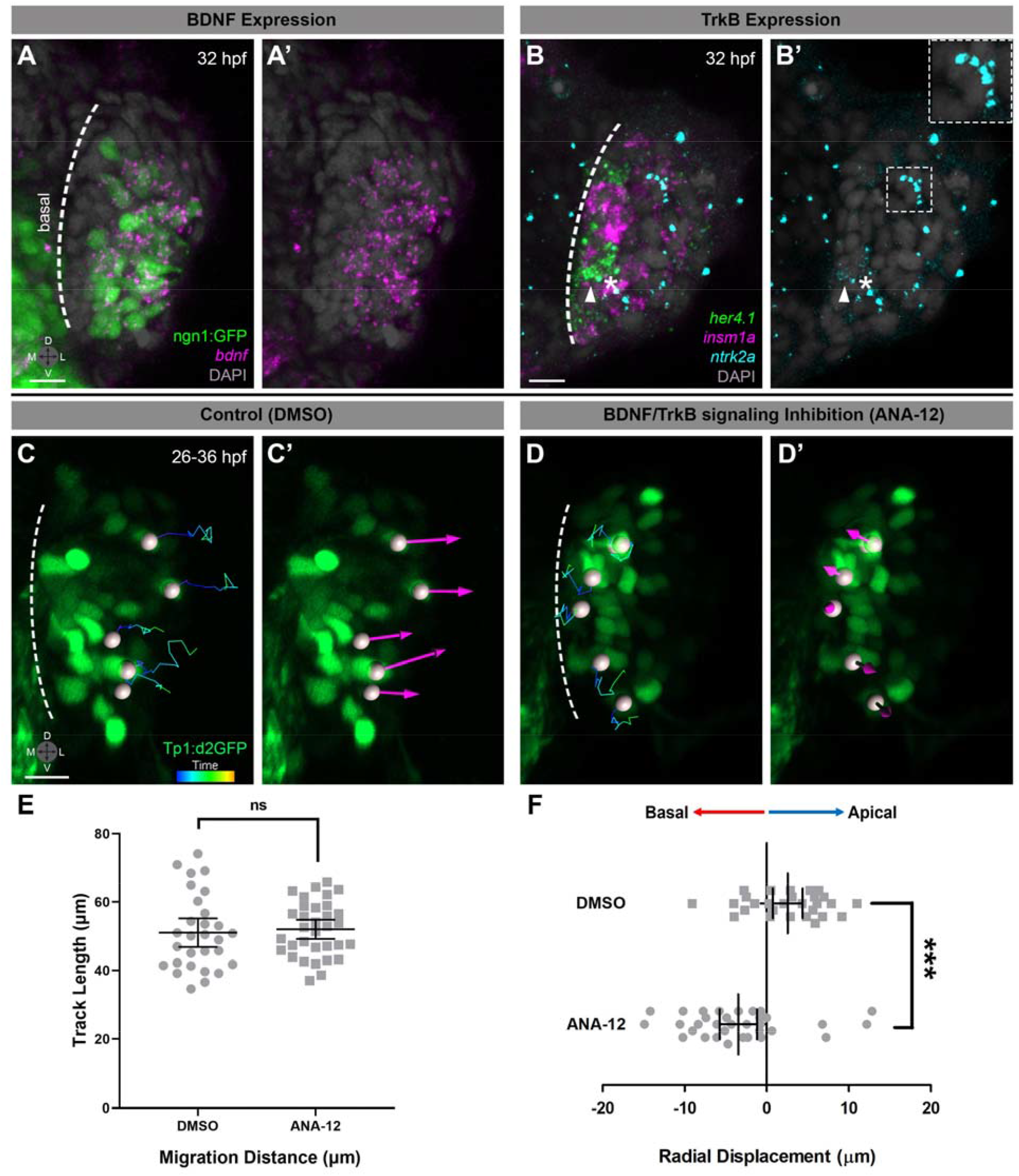
Neuronal BDNF facilitates basal to apical migration of progenitors via TrkB signaling. (A-A’) Optical section (z = 16 µm) of a representative ngn1:GFP+ embryo fixed at 32 hpf, followed by wmHCR targeting *bdnf* mRNA. (B-B’) Optical section (z = 6 µm) of a representative wild-type embryo fixed at 32 hpf, followed by wmHCR targeting *her4.1, insm1a*, and *ntrk2a* mRNA. An example neighborhood is indicated with a center cell (arrowhead) and contacting neighbor cell (asterisk) both expressing *ntrk2a*. Another medially-located cell, likely migrating apically, expresses *ntrk2a* (B’, box). (C-D’) Cell migration tracks (C,D) and displacement arrows (C’,D’) demonstrate movement of Tp1:d2GFP+ cells in embryos treated with DMSO (Control; C-C’) or 10 µM ANA-12/DMSO (BDNF-TrkB signaling inhibition; D-D’) from 22 hpf and imaged every 30 mins at 26-36 hpf. Dashed lines in (A-D’) indicate the OE’s basal edge. (E) Migration distance and radial displacement (F) of Tp1:d2GFP+ cells over 8 hours of tracking. Number of cells: DMSO (n = 28 from 7 embryos); ANA-12 (n = 33 from 8 embryos). Error bars: mean and 95% confidence intervals. Statistical significance: ****p < 0.0001. ngn1:GFP: green; bdnf: magenta; her4.*1*: green; *insm1a*: magenta; *ntrk2a*: cyan; DAPI: grey; Tp1:d2GFP: green. Scale bars: 10 µm.

### Neurogenic neighborhoods assemble and expand to facilitate olfactory regeneration

Given the constant turnover of OSNs beyond developmental time points, we investigated what role neurogenic neighborhoods might play in olfactory neuroregeneration. We induced OSN damage and death with copper sulphate^39, 40^ at 120-124 hpf, when developmental olfactory neurogenesis has almost completely ceased, and found that at 124 hpf, the number of basally-located neighborhoods had not changed (Fig **7C**), but the number of cells per neighborhood had rapidly increased (Fig **7A-B****”**, arrowheads and asterisks, supplementary videos 5 & 6). We found no difference in the number of center or direct neighbor cells (Fig **7D**); instead, neighborhood expansion was driven by a significant increase in the number of ‘indirect neighbor’ cells, which exhibit *her4.1*^*LOW*^ *insm1a*^*HIGH*^ expression and are contiguous with direct neighbor cells (Fig **7A-D**, hashtags, supplementary videos 5 & 6). Thus, injury-responsive olfactory regeneration triggers an immediate, accelerated commitment of existing neighborhoods towards a neuronal state.

**Figure 7:**
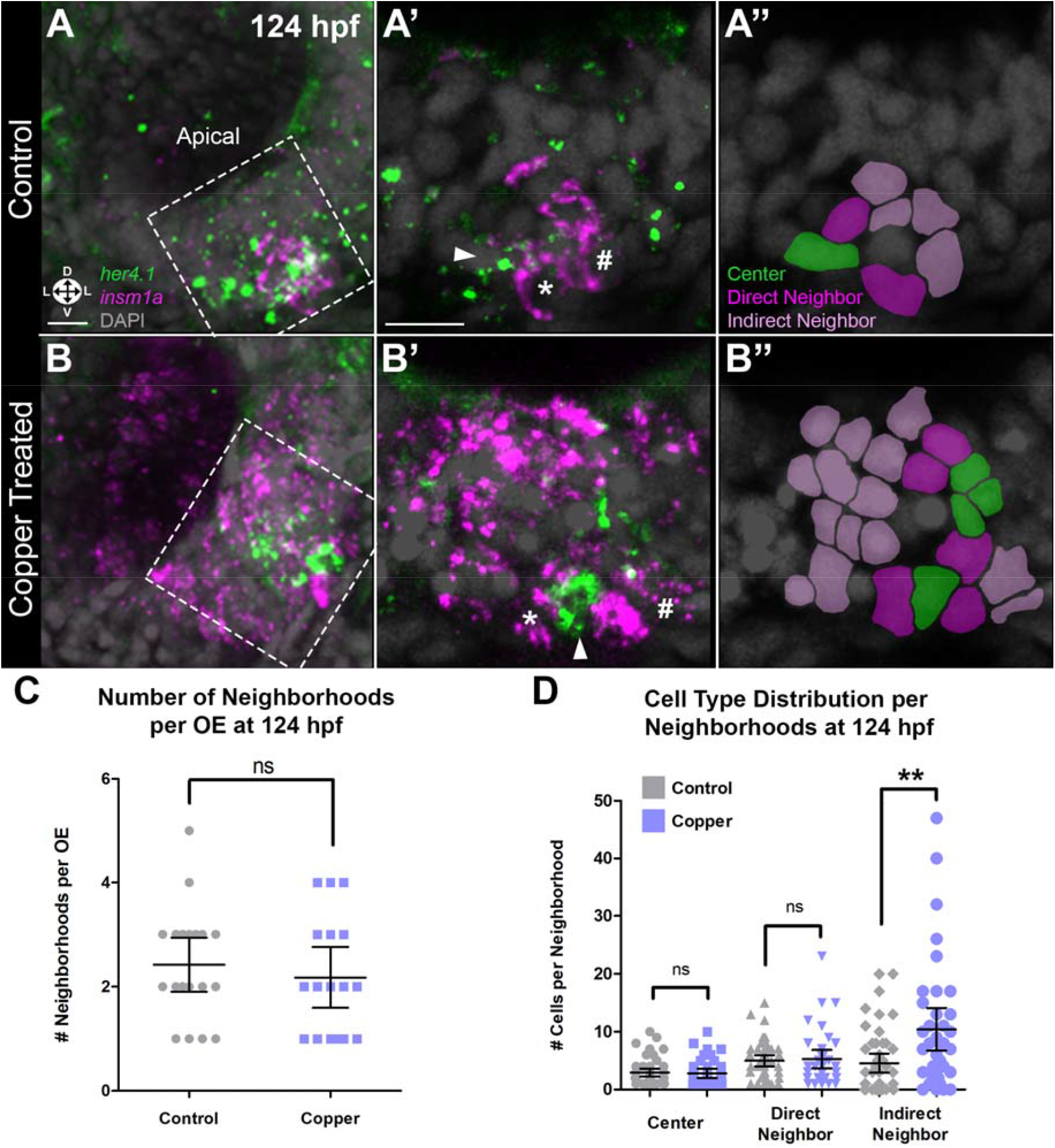
Neurogenic neighborhoods expand rapidly prior to olfactory regeneration. (A-B’’) 3D projections (A,B) and optical sections (A’-A’’,B’-B’’; z = 4 µm, boxed regions from A,B) of OEs after egg water (A-A’’) or 10 μM CuSO_4_/egg water (B-B’’) treatment from 120 hpf to 124 hpf, followed by fixation at 124 hpf and wmHCR for *her4.1* and *insm1a* mRNA. Arrowhead: center cell; asterisk: direct neighbor cell; hashtag: indirect neighbor cell. Schematics in (A’’,B’’) indicate center cells (green), direct neighbor cells (dark magenta), and indirect neighbor cells (light magenta). (C) Number of neighborhoods per OE at 124 hpf. Number of larvae: Control (n = 19); Copper (n = 17). (D) Number of center, direct neighbor, and indirect neighbor cells per neighborhood at 124 hpf. Number of neighborhoods: Control (n = 46 from 19 larvae); Copper (n = 38 from 17 larvae). Horizontal bars: mean and 95% confidence intervals. Statistical significance: ns (not significant) p > 0.05, *p < 0.05, **p < 0.01. *her4.1*: green; *insm1a*: magenta. Scale bars: 10 µm.

## DISCUSSION

Our findings have uncovered how neighborhoods of progenitor cells in the basal OE convert stochastic extrinsic and intrinsic signals into neuronal fate commitment. In cells with high levels of Notch signaling and target gene *her4.1* expression, *insm1a* expression and proneural gene function are inhibited, whereas in cells with high levels of *insm1a*, Notch signaling is inhibited, in turn promoting proneural gene expression and subsequent neurogenesis. This mutual antagonism maintains the balance between a continued progenitor state or a differentiated neuronal state and is regulated by RA signaling from the adjacent developing eye. Probabilistic modeling and experimental validation reveal the presence of a multistable configuration in which a tightly regulated bistable toggle switch is fine-tuned by fluctuations in RA and Notch signaling and promotes the self-assembly of spatially-restricted neurogenic neighborhoods. Newly differentiating OSNs emerge from neighborhoods in response to a BDNF-TrkB signaling gradient and migrate towards apically-located mature OSNs. Finally, expansion of neurogenic neighborhoods precedes and likely drives OSN regeneration. Taken together, our findings provide a multiscale, holistic view of how stochastic signaling inputs within single cells, in groups of cells, and within/across organs work together to develop and sustain a complex neuronal structure *in* vivo (Fig **8**).

**Figure 8:**
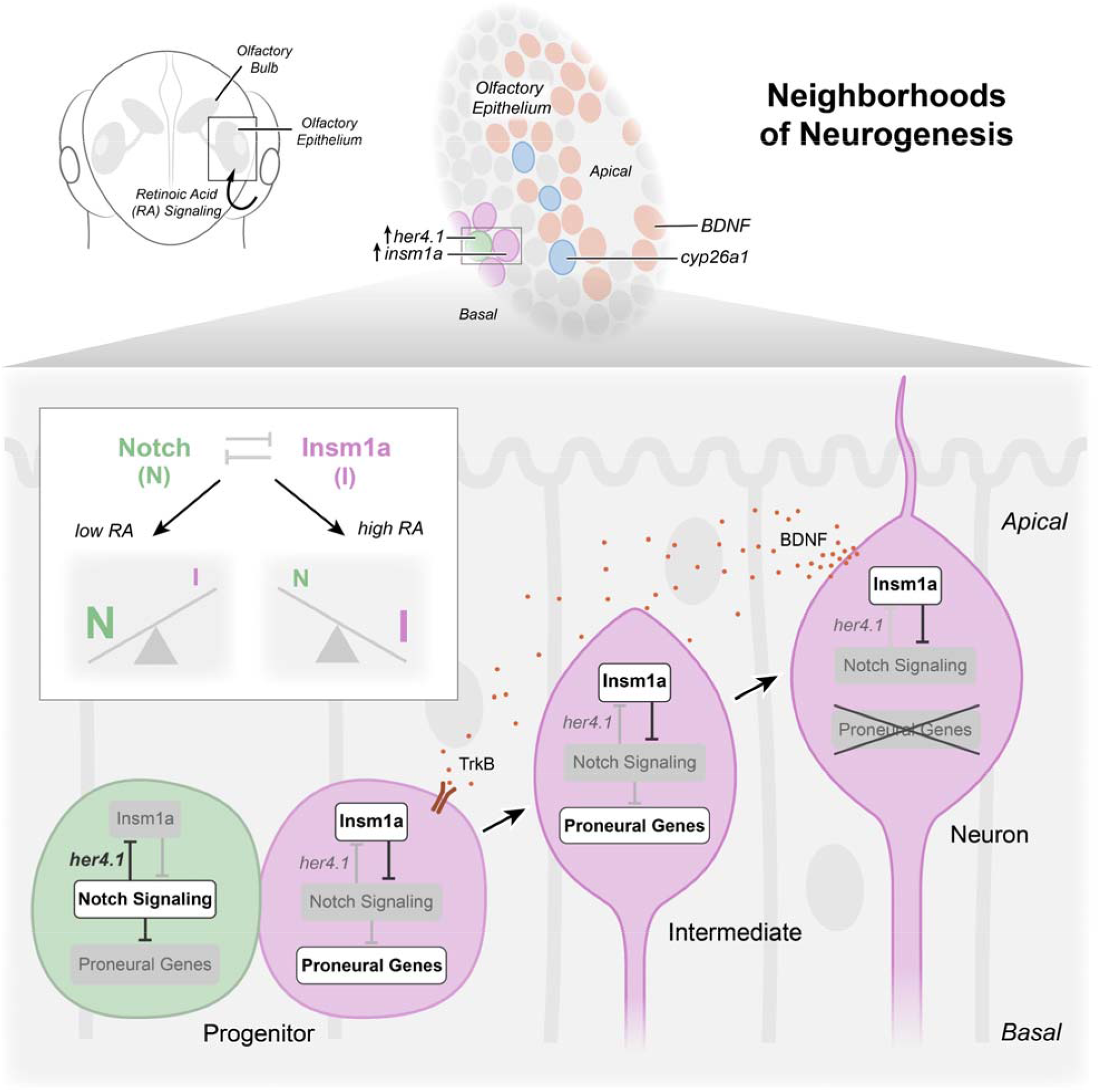
Overview of mechanisms driving sustained olfactory neurogenesis.

In many developing neuronal systems, progenitors have been suggested to adapt to substantial cell-to-cell variation and stochasticity by altering modes of cell division and differentiation^6, 41^ and, in some cases, participating in Notch signaling-dependent feedback^42^. Our stochastic modeling framework elucidates transition points between cell states in differentiating progenitors and predicts dynamic changes in system states during the process of continuous olfactory neurogenesis. In doing so, we uncover a previously unknown paradigm of cellular neighborhood self-assembly through which the OE integrates fluctuating and stochastic signaling inputs to streamline dynamic fate commitment, differentiation, and integration into the olfactory neuronal rosette. This neighborhood self-assembly is reminiscent of previous work in *Drosophila* demonstrating Notch signaling’s contribution to the self-organization of proneural clusters at sites of sensory hair cell differentiation^29, 43^. However, the vertebrate olfactory neighborhoods described here are highly dynamic and much less stereotyped, likely due to the stochastic interplay between inter-organ RA signaling and intrinsic Notch/Insm1a signaling. Intriguingly, Notch signaling has been reported to regulate *insm1a* expression either negatively or positively during retinal development^21^ or retinal regeneration^44^, respectively, suggesting that Notch signaling and Insm1a exhibit varying, context-dependent interactions. Our work, meanwhile, identifies a unique, mutually inhibiting bistable toggle switch system between Notch signaling and Insm1a at single-cell scale and within a highly constrained group of cells. This arrangement gives rise to emergent cell fate decisions regulating tissue-wide neurogenesis in the context of both the developing and regenerative OE.

The diffusion of RA into adjacent tissues has been previously identified as important for the development of a variety of organ systems^45-47^, and we recently demonstrated that RA can promote the differentiation of injected cancer cells that migrate in between the developing eye and nose^38^. The expression of RA synthesizing enzymes in the developing eye, consistent with previously described patterns^47, 48^, and the striking absence of those enzymes in the OE during time points of peak olfactory neurogenesis together point to an inter-organ signaling mechanism. Our targeted manipulations of RA levels suggest that RA production and diffusion from the eye acts as a critical pro-neurogenic influence on the composition and function of neurogenic neighborhoods. Interestingly, mechanical interactions between the developing eye and OE were recently shown to affect OE morphology^49^, which, together with our results, indicate significant mechanochemical interplay between the developing eye and nose.

Sharp changes in Notch and RA signaling have been previously shown to attenuate noise in signal transduction resulting from stochastic variation and fluctuations in gene expression across space and time^4, 5, 28^. Consistent with those findings, our work identifies spatially-conserved, *cyp26a1*-expressing cells that are located apical to neighborhoods of differential Notch signaling and likely control local RA availability to divide the developing OE into two zones – neighborhoods of neurogenesis in the basal region and fully differentiated neurons in the apical region. Thus, fluctuations in RA signaling from the eye may act as a source of stochasticity that alters the balance of *her4.1* and *insm1a* expression and is interpreted by the bistable toggle switch to shift neighborhood composition. This feedback mechanism produces stark differences between center and neighbor cells that drive RA-regulated neighborhoods to continuously promote neuronal differentiation.

Neurogenesis in complex tissues requires tight coordination of the acquisition of cell fate and the migration of newly formed neurons to their terminal locations^50, 51^. A role for BDNF signaling in neurogenesis has been controversial in the olfactory system, with some studies claiming no expression or requirement for BDNF and/or TrkB during olfactory development^52, 53^, while yet others have shown BDNF expression in the mature OE^54-56^. Here, consistent with *in vitro* studies of cultured human olfactory progenitor cells^37^, we reveal a key *in vivo* role for BDNF-TrkB signaling in the directed, basal-to-apical migration of new neurons exiting from neighborhoods, ensuring the arrival of a steady stream of functional OSNs at the olfactory surface.

Olfactory functionality, i.e. the ability to smell, is critical to an organism’s survival, and thus olfactory regeneration must proceed efficiently after injury. Consistent with previous descriptions of active Notch signaling in stem cells in the lesioned murine adult OE^14, 17, 57^, our experiments demonstrate that basal progenitors in the regenerating zebrafish OE rapidly redeploy bistable switch-driven neighborhoods of neurogenesis. Given the sequence of events known to be important for neuroregenerative recovery and repair^58, 59^, we had anticipated that copper-induced olfactory injury would result in a lag time prior to the initiation of neurogenesis, but to our surprise, we found an immediate expansion of neighborhoods, suggesting that their ordering and growth is one of the earliest components of a post-injury neuroregenerative response. Expansion of existing neighborhoods rather than *a de novo* increase in their number suggests that regeneration is triggered using already present multicellular patterning that may be constrained by the contact-dependent nature of Notch signaling. Moving forward, targeted injury of the OE will prove useful for investigating the plasticity of progenitor cells within the neighborhood architecture and how they respond *in vivo* to a variety of regenerative stimuli.

Our findings highlight the diversity of cell biological events that contribute to the generation and assembly of functional neurons. While the specification of cell fate is clearly a critical aspect of neurogenesis, parallel processes such as the earlier self-assembly of dynamic progenitor cell groups and the ultimate movement of maturing neurons to their final destinations are equally important for the formation of an intricate sensory organ. Thus, multicellular self-organization can serve as a fundamental paradigm for integrating stochastic cell-cell signaling to streamline continuous neurogenesis *in vivo*.

## ACKNOWLEDGMENTS

We are grateful to Katie Harvey and Dr. Ana Beiriger for assistance with figure preparation and creating the model (Fig. 8); Dr. Ana Beiriger, Dr. Thomas Bozza, Dr. Elizabeth LeClair, and Dr. Abigail Green-Saxena for feedback on the manuscript; Nathan Burg for zebrafish husbandry assistance; and all Saxena Lab members for feedback on the project. We thank Dr. Yoshihiro Yoshihara, RIKEN BSI, and National Bioresource Project of Japan for *Tg(OMP2k:lyn-mRFP)/rw03*; Dr. Robert Kelsh for *Tg(−4.9sox10:eGFP)*; Dr. Caroline Burns and Dr. Geoff Burns for *Tg(hsp70l:DN-MAML-GFP)* and *Tg(hsp70l:1xMYC-notch1a-intra)*; Dr. Brian Link for *Tg(Tp1bglob:d2GFP)*; Dr. Uwe Strahle for *Tg(−8.4neurog1:nRFP)* and *Tg(−8.4neurog1:GFP)*; and Dr. Ann Morris for *insm1a* promoter plasmids. Funding for this work was provided by National Institutes of Health R01HD100023; a Pilot Project Award from the NSF-Simons Center for Quantitative Biology at Northwestern University, an NSF-Simons MathBioSys Research Center (supported by a grant from the Simons Foundation/SFARI (597491-RWC) and the National Science Foundation (1764421)); and the Chicago Biomedical Consortium with support from the Searle Funds at The Chicago Community Trust (all to A.S.). J.G. was supported in part by HSI-STEM grant #P031C160237 from USA ED.

## AUTHOR CONTRIBUTIONS

Conceptualization, S.G.R., J.N.L., J.L., and A.S.; methodology, S.G.R., J.N.L., L.M.N., F.M., and A.S.; formal analysis, S.G.R., J.N.L., F.M., and K.W.; investigation, S.G.R., J.N.L., L.M.N., F.M., K.W., and J.G.; resources, L.M.N., J.L., and A.S.; writing – original draft, S.G.R. and A.S.; writing – review and editing, S.G.R., J.N.L., F.M., J.L., and A.S.; visualization, S.G.R., J.N.L., F.M., K.W., and A.S.; supervision, J.L. and A.S.; funding acquisition, A.S.

## Supplementary Figures

**Figure S1:**
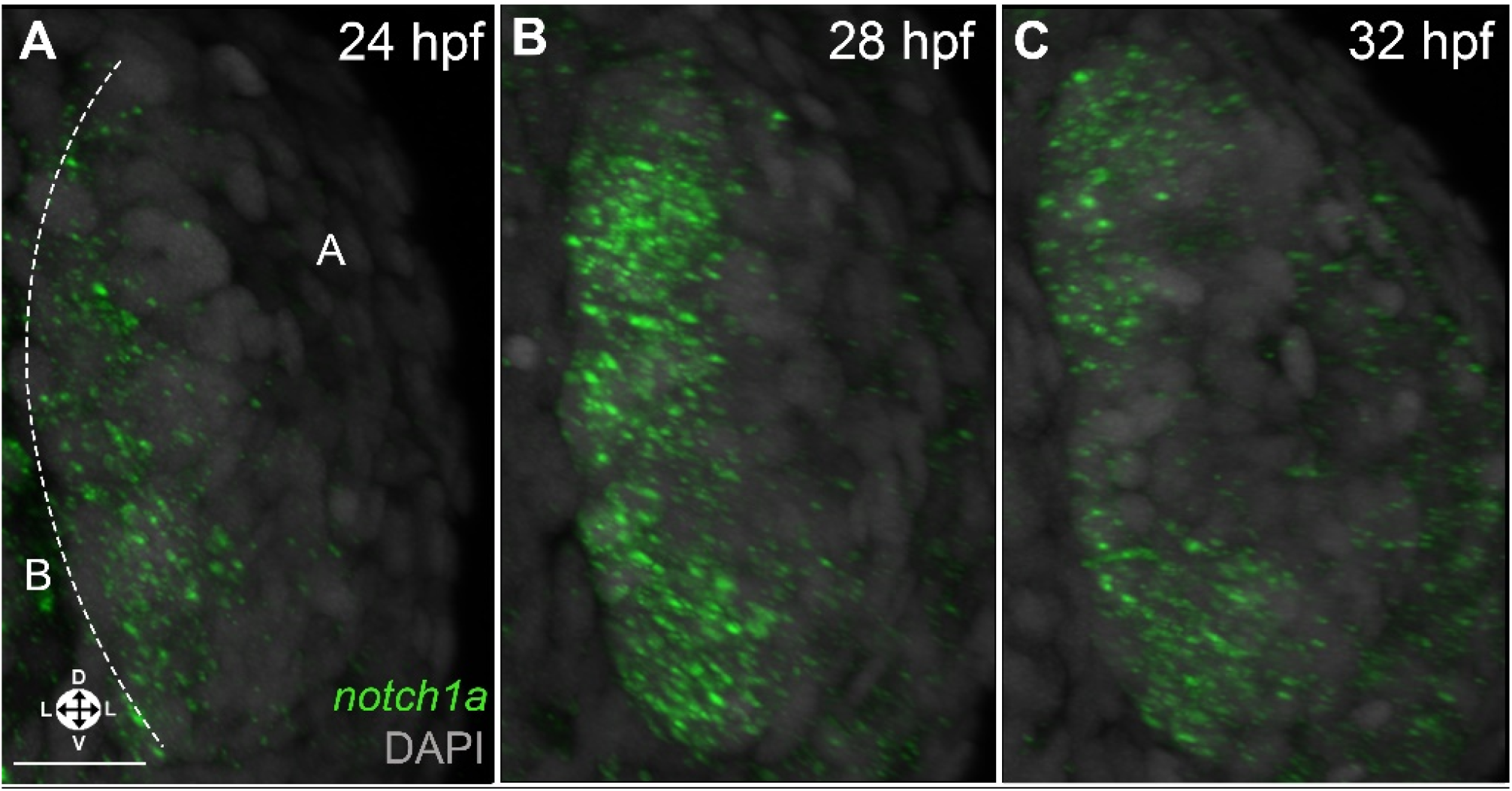
*notch1a* mRNA is expressed in the basal OE. (A-C) 3D projection anterior views of wmHCR of OEs at 24 hpf (A), 28 hpf (B), and 32 hpf (C) showing basal expression of *notch1a* mRNA. n = 4 embryos per time point. *notch1a*: green; DAPI: grey. B: basal; A: apical. Orientation arrows: L: Lateral; D: dorsal; V: ventral. Scale bars: 20 µm.

**Figure S2:**
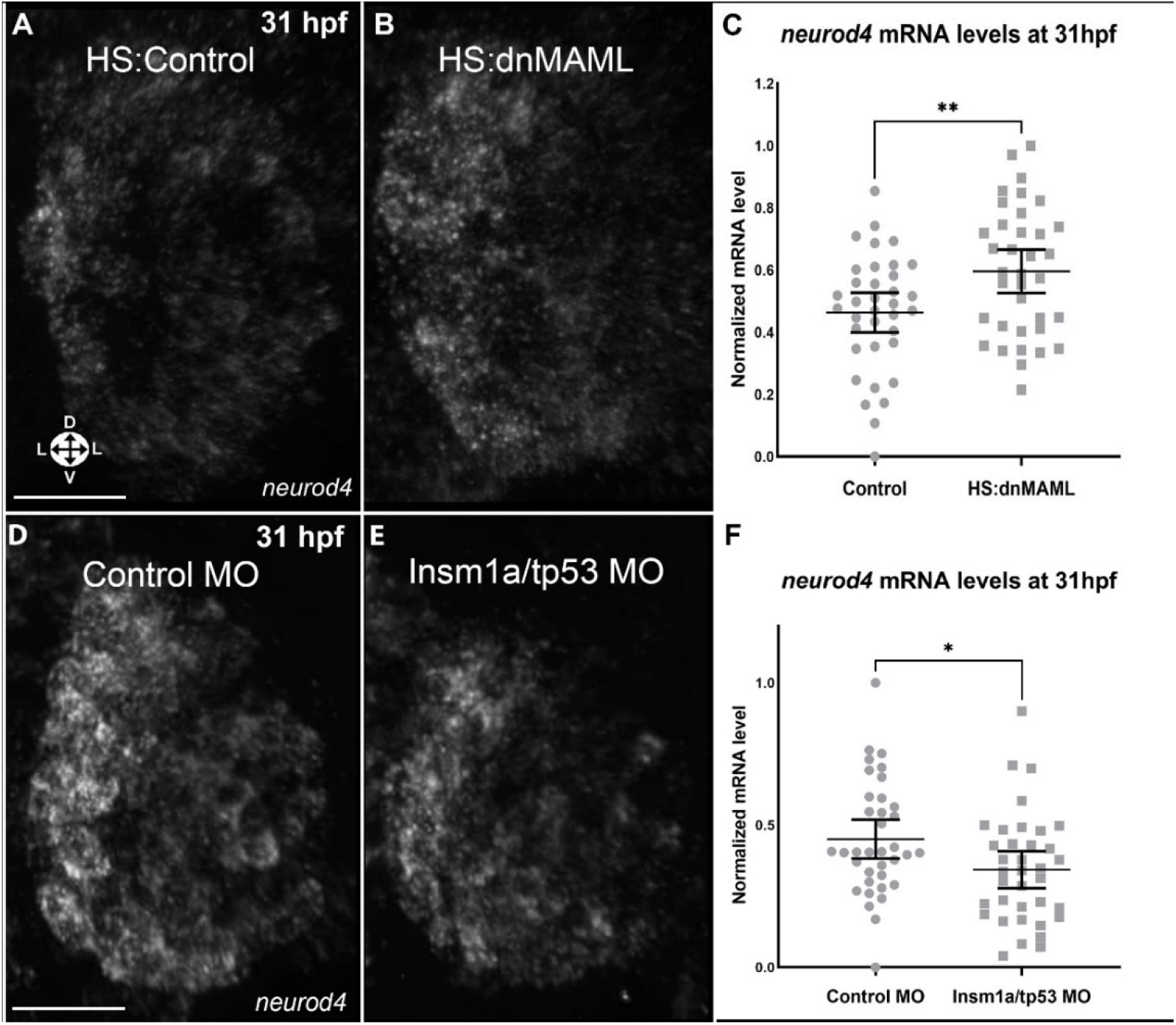
Notch signaling and Insm1a have opposing effects on *neurod4* expression. (A-B) 3D projection anterior views of OEs from embryos heat shocked at 24 hpf without (A) or with (B) the heat shock-inducible transgene HS:dnMAML, followed by fixation at 31 hpf and wmHCR for *neurod4* mRNA. (C) Normalized *neurod4* HCR signal intensity per cell. Number of cells: Control (n = 36 from 3 embryos); HS:dnMAML (n = 36 from 3 embryos). (D-E) 3D projection anterior views of OEs from embryos injected with control MO (D) or Insm1a/tp53 MO (E), followed by fixation at 31 hpf and wmHCR for *neurod4* mRNA. (F) Normalized *neurod4* HCR signal intensity per cell. Number of cells: Control MO (n = 36 from 6 embryos); Insm1a/tp53 MO (n = 36 from 6 embryos). Horizontal bars: mean and 95% confidence intervals. Statistical significance: *p < 0.05, **p < 0.01. *neurod4*: white. Orientation arrows: L: Lateral; D: dorsal; V: ventral. Scale bars: 30 µm.

**Figure S3:**
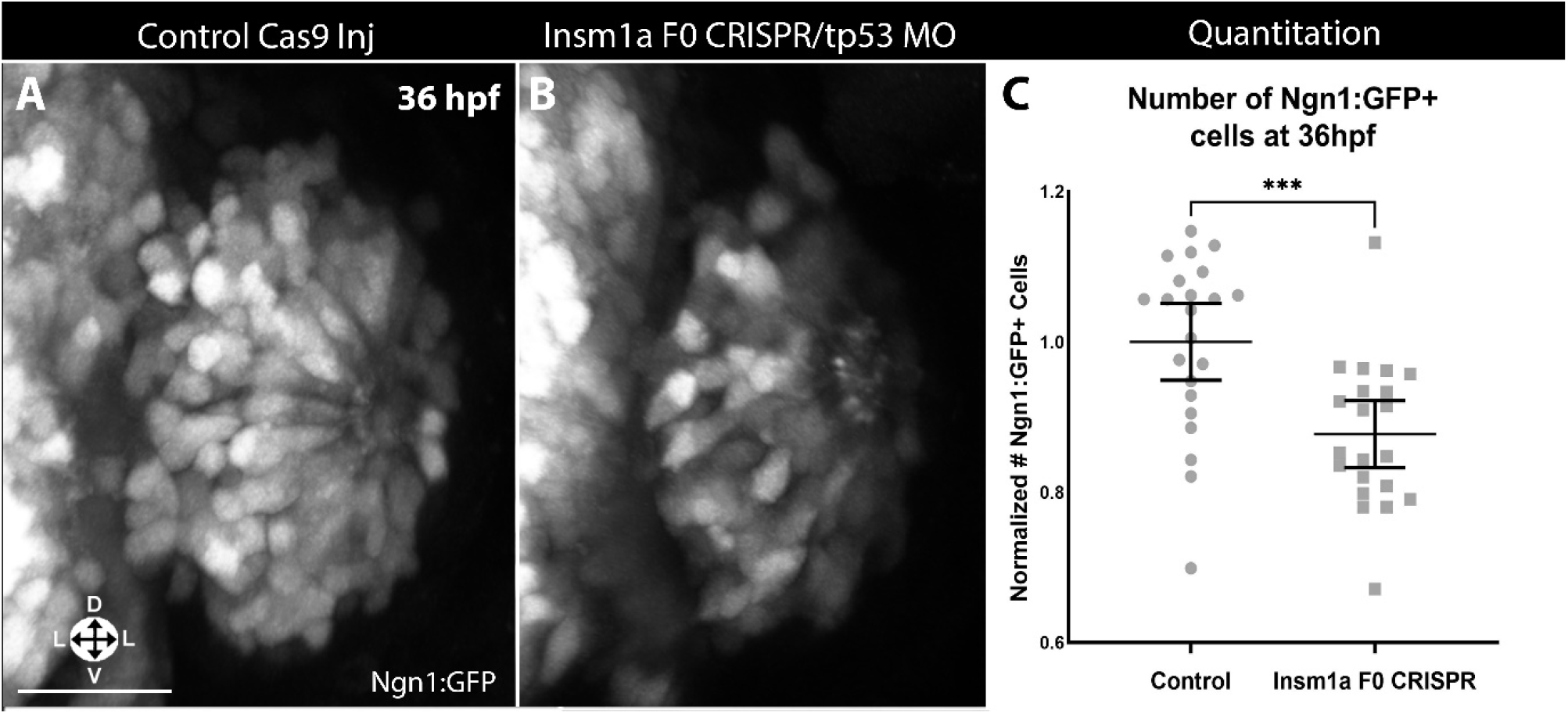
F_0_ CRISPR/Cas9 knockdown of Insm1a yields reduced neuron numbers. (A-B) 3D projection anterior views of OEs from Ngn1:GFP+ embryos injected with SpCas9-2NLS without (A) or with two redundant synthetic guide RNAs (sgRNAs) targeting *insm1a* along with tp53 MO (B), followed by fixation at 36 hpf. (C) Normalized number of GFP+ cells per embryo at 36 hpf. Control (n = 22 embryos); Insm1a F0 CRISPR (n = 21 embryos). Horizontal bars: mean and 95% confidence intervals. Statistical significance: ***p < 0.001. Ngn1:GFP: white. Orientation arrows: L: Lateral; D: dorsal; V: ventral. Scale bars: 30 µm.

**Figure S4:**
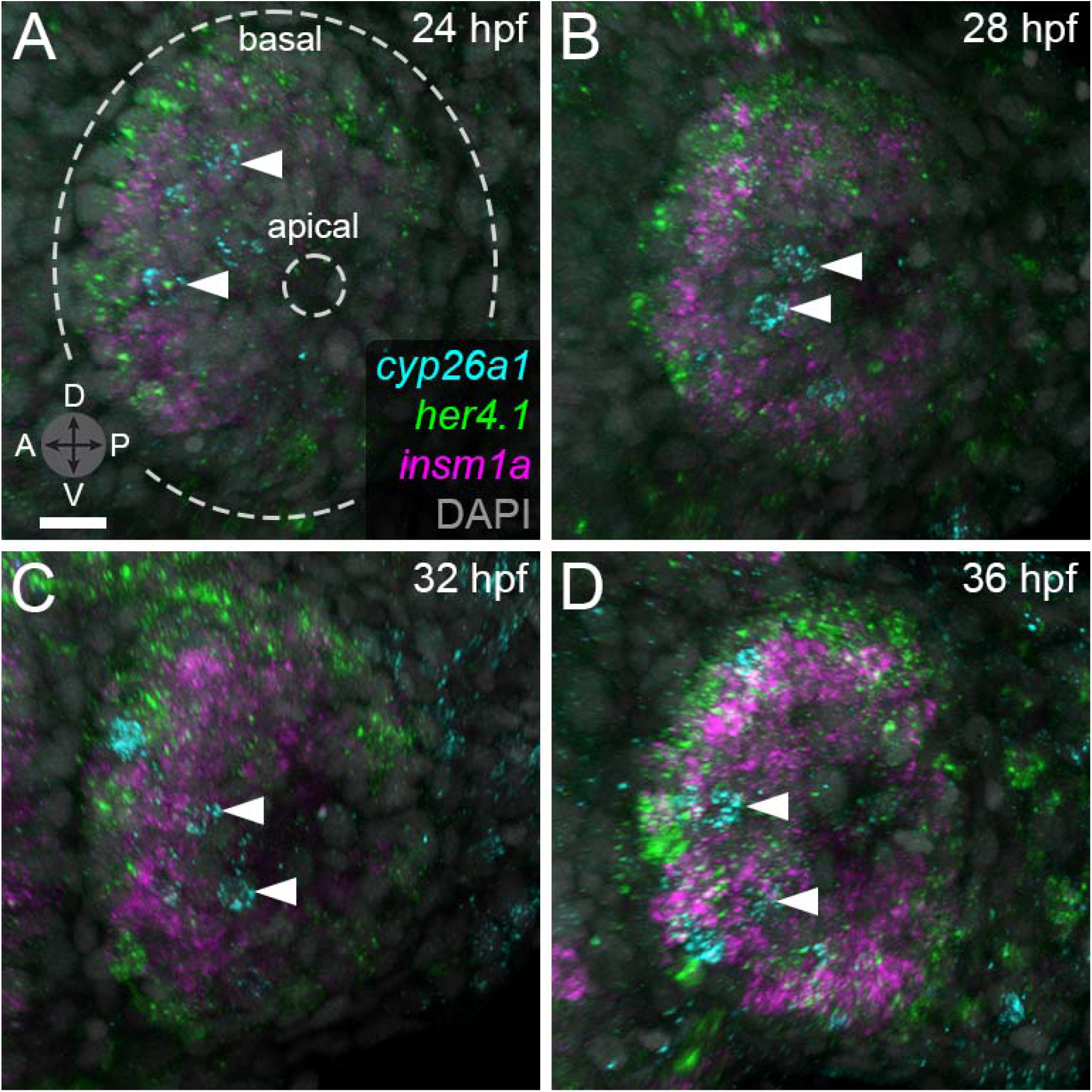
*cyp26a1*+ cells are located apically relative to neighborhoods. (A-D) 3D projection anterior views of OEs from representative embryos fixed at 24 hpf (A), 28 hpf (B), 32 hpf (C) and 36 hpf (D), followed by wmHCR targeting *cyp26a1, her4,1*, and *insm1a* mRNA. Dashed lines outline the OE, and arrowheads indicate *cyp26a1*+ cells located apical to *her4.1*/*insm1a* neighborhoods. n = 6 embryos per time point. *her4.1*: green; *insm1a*: magenta; *cyp26a1*: blue; DAPI: grey. Orientation arrows: D, dorsal; V, ventral; A, anterior; P, posterior. Scale bar: 30 µm.

**Figure S5:**
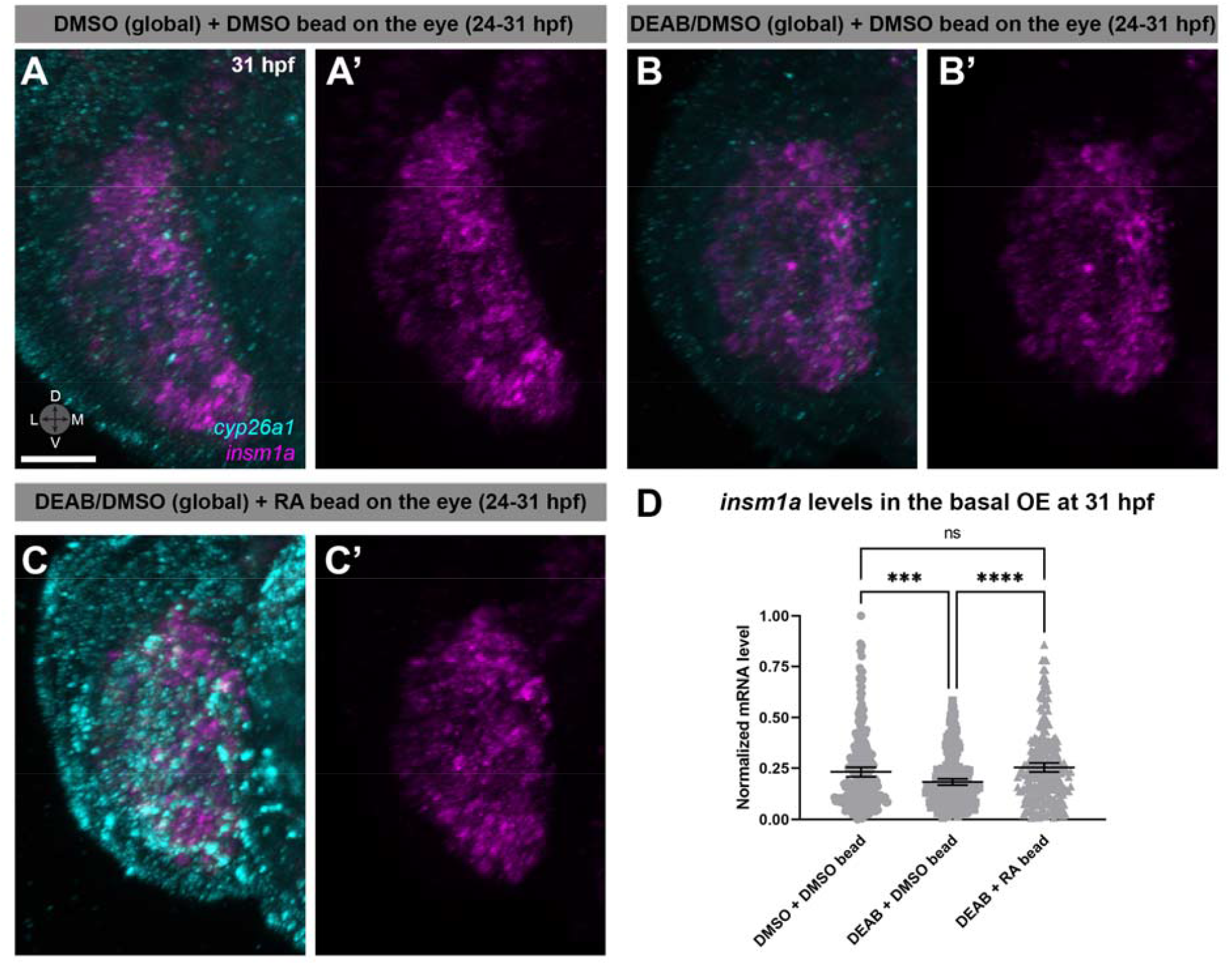
Overexpression of retinoic acid (RA) in the eye reverses effects of RA inhibition. (A-B’) 3D projections of representative OEs from embryos treated with DMSO (A,A’) or 10 µM DEAB/DMSO (B,B’) from 24 to 31 hpf, with a bead soaked in DMSO placed on the adjacent eye in both cases. (C,C’) 3D projections of representative OEs from embryos treated with 10 µM DEAB/DMSO from 24 to 31 hpf, with a bead soaked in 100 µM RA/DMSO placed on the adjacent eye. All embryos were fixed at 31 hpf, followed by wmHCR targeting *insm1a* and *cyp26a1* mRNA. (D) Normalized *insm1a* HCR signal intensity per cell. Number of cells: DMSO + DMSO bead (n = 257 from 6 embryos); DEAB + DMSO bead (n = 253 from 6 embryos); DEAB + RA bead (n = 248 from 6 embryos). Horizontal bars: mean and 95% confidence intervals. Statistical significance: ns (non-significant), ***p < 0.001, ****p < 0.0001. *cyp26a1*: cyan; *insm1a*: magenta. Orientation arrows: D, dorsal; V, ventral; M, medial; L, lateral. Scale bars: 20 µm.

**Figure S6:**
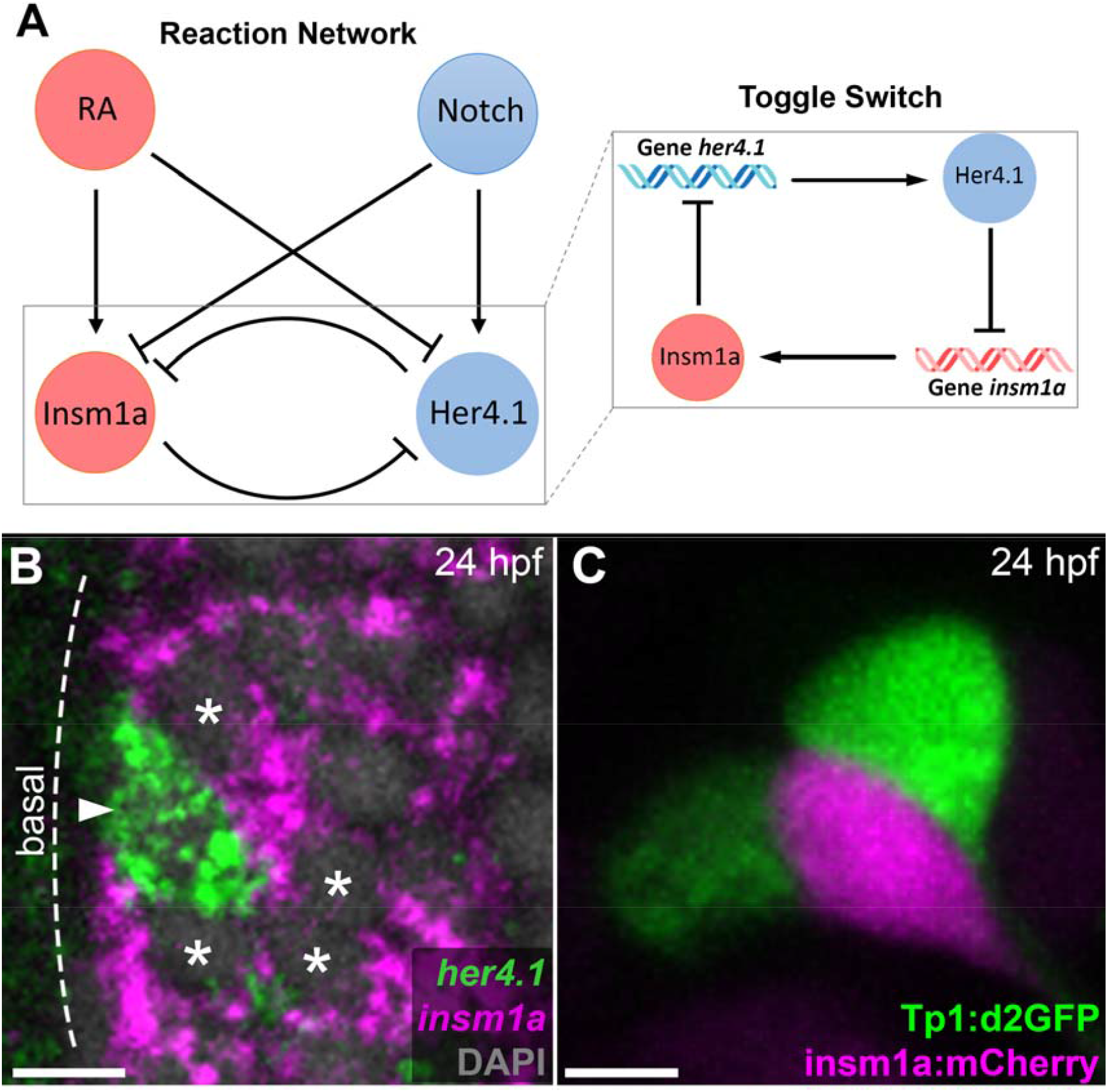
Reaction network contains a *her4.1*/*insm1a* bistable toggle switch. (A) Reaction network of an RA- and Notch signaling-regulated toggle switch system. (B) Optical section (z = 1.25 µm) of a representative wild-type embryo fixed at 24 hpf, followed by wmHCR targeting *her4.1* and *insm1a* mRNA. An example neighborhood is shown with a center cell (high *her4.1*, low *insm1a*; arrowhead) surrounded by contacting neighbor cells (low *her4.1*, high *insm1a*; asterisks). Dashed line indicates the OE’s basal edge. (C) Optical section (z = 10 µm) of a representative Tp1:d2GFP+; mosaic insm1a:Cherry+ embryo at 24 hpf recapitulating the data in (B) with reporter lines. *her4.1*: green; *insm1a*: magenta; DAPI: grey; Tp1:d2GFP: green; insm1a:mCherry: magenta. Scale bars: 10 µm.

**Figure S7:**
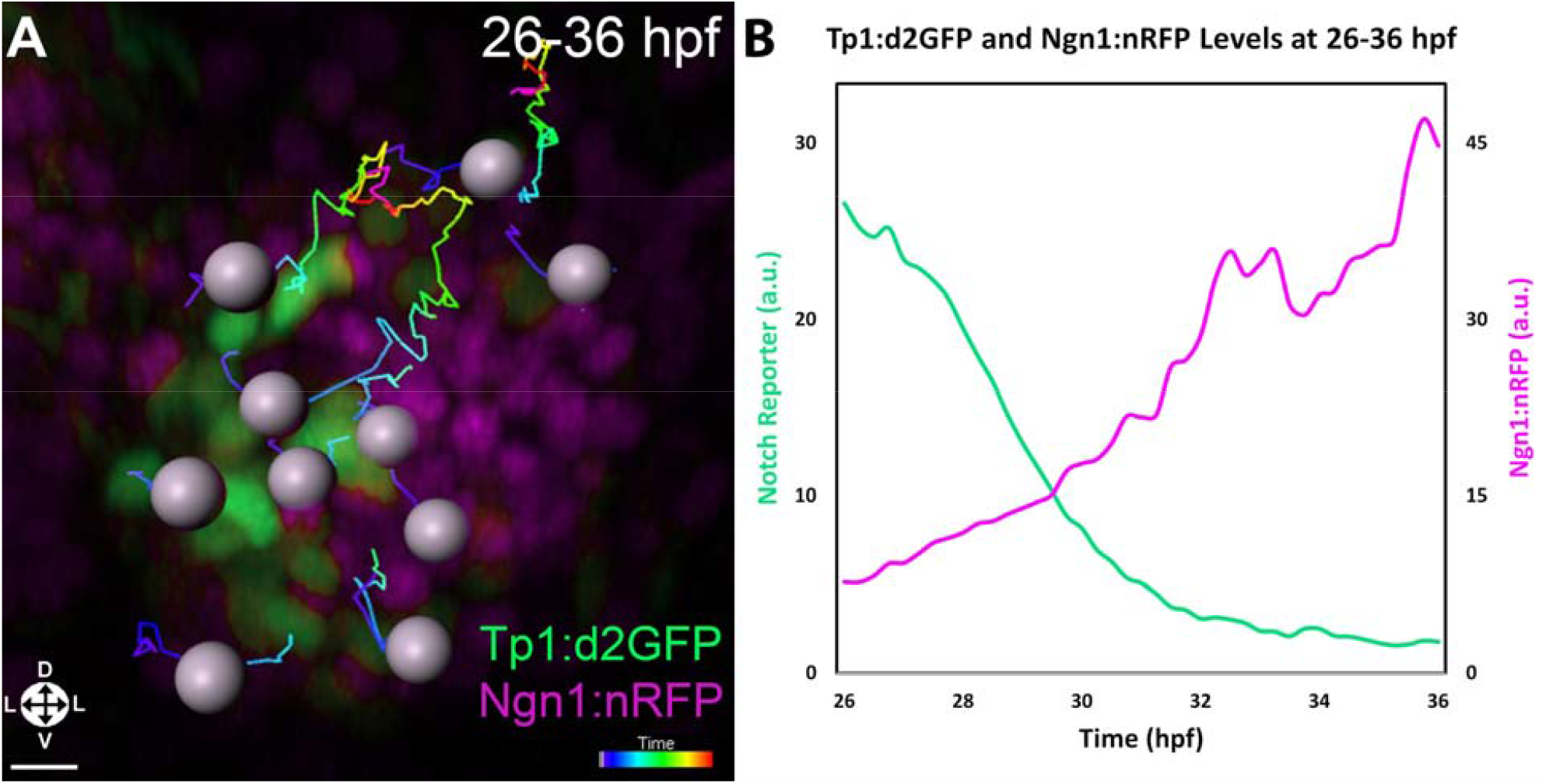
Notch signaling-positive center cells eventually differentiate into neurons. (A) 3D projection anterior view of an OE from a representative Tp1:d2GFP+; Ngn1:nRFP+ embryo imaged every 15 minutes at 26-36 hpf, followed by tracking of individual cells over time (cell tracks). (B) Quantitation of mean GFP (green line) and RFP (magenta line) signal intensities across individually tracked cells (n = 19 cells from 2 embryos). Tp1:d2GFP: green; Ngn1:nRFP: magenta. Orientation arrows: L: Lateral; D: dorsal; V: ventral. Scale bar: 15 µm.

**Figure S8:**
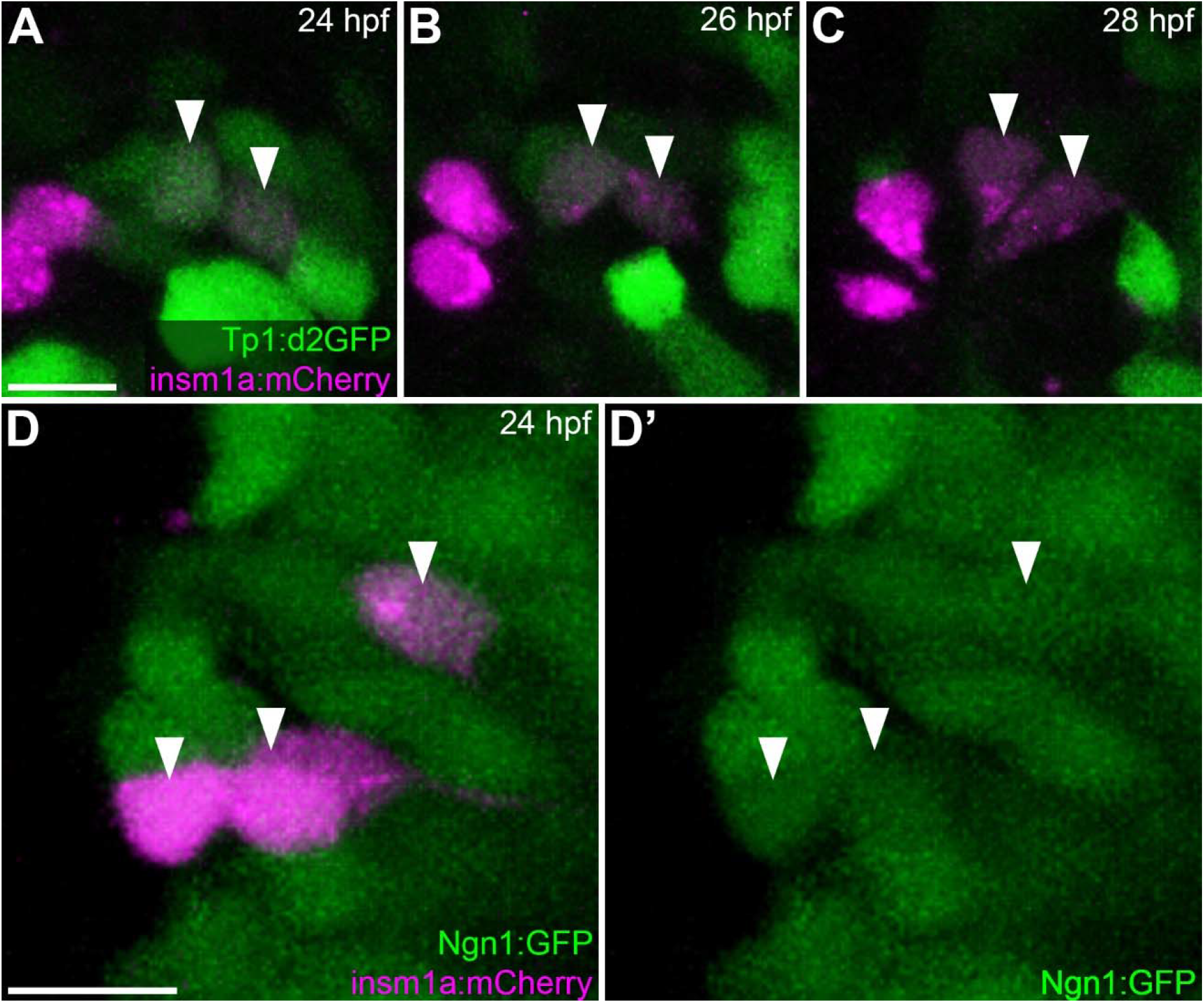
Center cells transition into neighbor cells that differentiate into neurons. (A-C) Optical sections (z = 15 µm) of the OE from a representative Tp1:d2GFP+; mosaic insm1a:mCherry+ embryo with d2GFP+ cells transitioning into mCherry+ cells (white arrowheads) at 24 hpf (A), 26 hpf (B), and 28 hpf (C). 8 transitioning cells were observed across 9 embryos. (D-D’) Optical section (z = 15 µm) of the OE from a representative Ngn1:GFP+; mosaic insm1a:mCherry+ embryo. Arrowheads indicate GFP+; mCherry+ cells. 64/72 insm1a:mCherry+ cells were also Ngn1:GFP+ across 9 embryos. Tp1:d2GFP: green; insm1a:mCherry: magenta; Ngn1:GFP: green. Scale bar: 10 µm.

**Figure S9:**
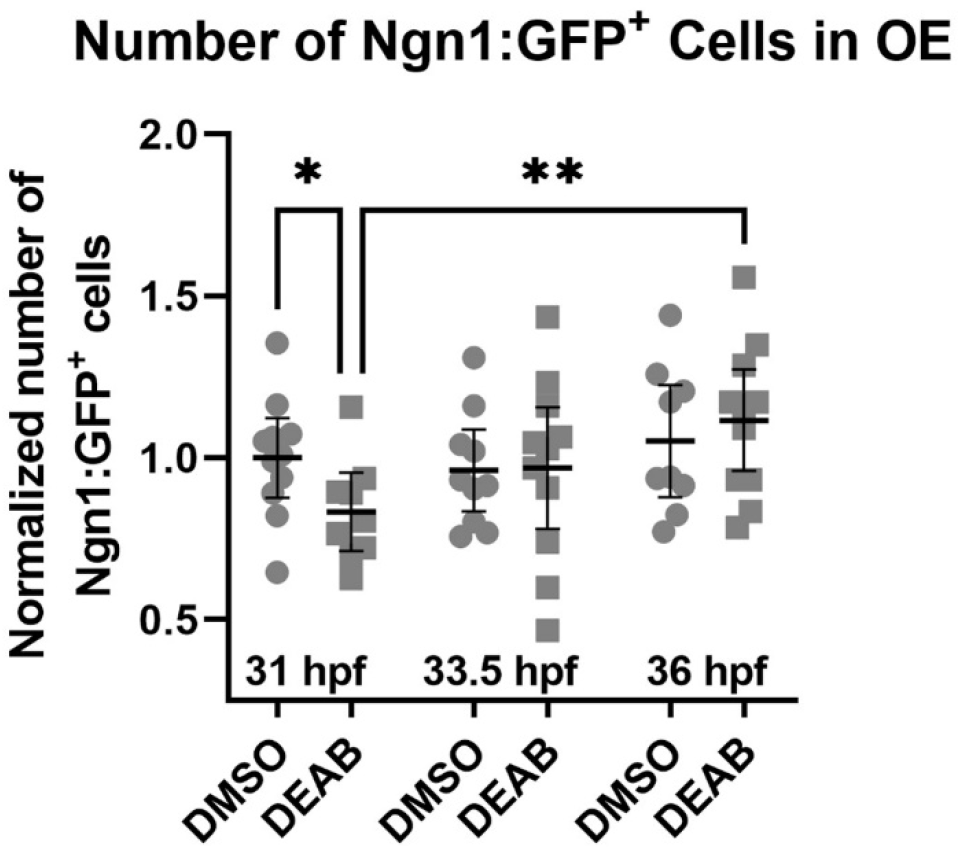
Attenuation of RA synthesis alters number of neurons. Mean**-**normalized number of Ngn1:GFP+ cells per embryo at 31 hpf, 33.5 hpf and 36 hpf after treatment with DMSO (negative control) or 10 µM DEAB/DMSO (RA inhibitor) from 24 to 31 hpf. n = 9 embryos per condition per time point.

## Supplemental Videos

**Supplementary Video 1. Tracking Notch signaling dynamics from 23 to 31 hpf**. Anterior view 3D projection of a time-lapse (23-31 hpf, 10 min. intervals) of transgenic Tp1:d2GFP+ embryos. Cell tracks label GFP+ basal cells migrating apically. Orientation: left: lateral; right: lateral; up: dorsal; down: ventral. Scale bar: 15 µm

**Supplementary Video 2. *cyp26a1* expression in control and RA-inhibited embryos**. 70 μm z-stack (z =1.25 μm optical sections) through OEs from wild-type embryos treated with DMSO (Control) or 10 µM DEAB/DMSO (RA synthesis inhibition) at 24-31 hpf, followed by wmHCR targeting *cyp26a1* mRNA. *cyp26a1*+ cells present in the medial OE exhibit decreased *cyp26a1* levels upon RA synthesis inhibition. *cyp26a1*: cyan; DAPI: grey. Orientation: left: lateral; right: lateral; up: dorsal; down: ventral. Scale bar: 10 μm

**Supplementary Video 3. Reaction network dynamics**. Time-evolving dynamics of the probability landscape paused at 24, 28, 32, and 36 hpf.

**Supplementary Video 4. *cyp26a1* expression in control and RA-inhibited embryos**. Anterior view 3D projections of OEs during time-lapse imaging (26-36 hpf, 10 min. intervals) of transgenic Tp1:d2GFP+ embryos. Cell migration tracks demonstrate movement of GFP+ cells in embryos treated with DMSO (Control) or 10 µM ANA-12/DMSO (BDNF-TrkB signaling inhibition) from 22 hpf and imaged every 30 mins at 26-36 hpf. Orientation: left: medial; right: lateral; up: dorsal; down: ventral. Scale bar: 10 µm

**Supplementary Video 5. Neighborhood composition in control and copper-treated larvae**. 3D projections of OEs from larvae treated with egg water (Control) or 10 μM CuSO_4_/egg water solution at 120-124 hpf, followed by wmHCR targeting *her4.1* and *insm1a*. Colored spots indicate center cells (green), direct neighbor cells (dark magenta), and indirect neighbor cells (light magenta) in neighborhoods. DAPI: grey. Orientation: left: lateral; right: lateral; up: dorsal; down: ventral. Scale bar: 10µm.

**Supplementary Video 6. Neighborhood composition in control and copper-treated larvae**. 24 um z-stack (z = 2.00 um optical sections) through OEs from wild-type larvae treated with egg water (Control) or 10 µM CuSO_4_/egg water solution at 120-124 hpf, followed by wmHCR targeting *her4.1* and *insm1a*. Colored circles indicate center cells (green), direct neighbor cells (dark magenta), and indirect neighbor cells (light magenta) in neighborhoods. DAPI: grey. Orientation: left: lateral; right: medial; up: dorsal; down: ventral. Scale bar: 10 µm

## MATERIALS AND METHODS

### RESOURCE AVAILABILITY

#### Lead Contact

Further information and requests for resources and reagents can be directed to the lead contact, Ankur Saxena.

#### Materials Availability

All unique reagents generated in this study are available from the Lead Contact.

#### Data and Code Availability

The data reported in this study are available from the Lead Contact.

### EXPERIMENTAL MODEL AND SUBJECT DETAILS

#### Zebrafish Model

AB Zebrafish were treated and cared for in accordance with the National Institutes of Health Guide for the Care and Use of Laboratory animals. All experiments were approved by the University of Illinois at Chicago Institutional Animal Care Committee. Zebrafish were housed in a secure aquatic facility in 3.5 L tanks with 10-20 adults per tank, daily health monitoring, and a diet of Skretting’s GEMMA Micro 75, 150, or 300, depending on age/size. Zebrafish embryos and larvae were grown, staged, and harvested as previously described^60, 61^ in egg water (15 mM NaCl, 8.3 mM CaSO_4_, methylene blue) at 28.5°C. Embryos/larvae at time points older than 24 hours post-fertilization (hpf) were treated with 1-phenyl-2-thiourea as needed to prevent pigmentation from interfering with imaging. Transgenic lines used and their abbreviations in this manuscript are: *Tg(Tp1bglob:d2GFP)*^23, 62^ = Tp1:d2GFP; *Tg(OMP2k:lyn-mRFP)/rw035*^63^ = OMP:RFP; *Tg(−4.9sox10:eGFP)*^64^ = Sox10:eGFP; *Tg(−8.4neurog1:nRFP)*^65^ = Ngn1:nRFP; *Tg(−8.4neurog1:GFP)*^65^ = Ngn1:GFP; *Tg(hsp70l:DN-MAML-GFP)*^16^ = HS:dnMAML. Insm1a:mCherry construct was assembled based on a previously described 2.4 kb *insm1a* promotor/enhancer region^21^, which we further modified with the Tol2kit system^66^ to create a 2.5 kb promoter/enhancer-driven reporter construct that drives mCherry expression. This construct was injected at the one-cell stage to generate mosaic F_0_ embryos for downstream imaging and analysis.

### METHOD DETAILS

#### Whole mount in situ hybridization chain reaction (wmHCR)

Zebrafish embryos and larvae were fixed using 4% (w/v) PFA in DEPC-treated PBS. Hybridization chain reaction (HCR) probes were designed to detect relative mRNA levels of *her4.1, insm1a, neurod4, ascl1a, aldh1a2, aldh1a3, cyp26a1, b-actin, bdnf*, and *ntrk2a*. HCR was performed as previously described^24^, and samples were counterstained with DAPI. All probes, fluorescent hairpins, and buffers were purchased from Molecular Instruments, Inc. (www.molecularinstruments.com).

#### Notch signaling knockdown and overactivation with heat shock

To inhibit Notch signaling, heat shock was induced at 24 hpf by incubating *Tg(hsp70l:DN-MAML-GFP)* heterozygotes or wild-type clutchmates at 37°C for 60 minutes, followed by 5 minutes at 40°C. Embryos were then sorted for the presence or absence of global dnMAML-GFP fluorescence before fixation at 31 hpf or 36 hpf.

#### Chemical treatments

For knockdown of retinoic acid (RA) synthesis, zebrafish clutchmates were incubated in the dark from 24 hpf to 31 hpf in 10 μM DEAB + 1% DMSO in egg water or 1% DMSO only in egg water, followed by either fixation at 31 hpf or washing of embryos with egg water and fixation at 33.5 hpf or 36 hpf. To rescue effects of RA inhibition, beads were soaked in 1% DMSO or 100 μM RA and placed on the left eye of embryos that are incubated in either 10 μM DEAB + 1% DMSO in egg water or 1% DMSO only in egg water underneath an agarose bed from 24 to 31 hpf. For BDNF-TrkB signaling inhibition, zebrafish clutchmates were incubated in the dark starting at 22 hpf in 10 μM ANA-12 + 1% DMSO in egg water or 1% DMSO only in egg water, followed by transfer at 26 hpf to agarose-based imaging molds (previously described^11, 67^) for time-lapse imaging. For copper treatments, zebrafish larvae were incubated in 10 μM CuSO_4_ solution in egg water from 120 to 124 hpf and fixed at 124 hpf.

#### Insm1a knockdown

Morpholinos were obtained from Gene Tools, LLC. As previously described^21^, a translation-blocking morpholino against Insm1a (GGTTGAAATCAGAGGCACACCT) was injected into one-cell stage embryos at 6 ng/embryo along with a tp53 morpholino (GCGCCATTGCTTTGCAAGAATTG) at 9 ng/embryo to suppress non-specific cell death. Clutchmates were injected with a standard control morpholino targeting a mutant variant of human β-globin (CCTCTTACCTCAGTTACAATTTATA).

For F_0_ CRISPR/Cas9 knockdown, we first established knockdown efficiency by testing control F_0_ CRISPR/Cas9 injections that targeted the gene *tyrosinase* (tyr) (synthetic guide RNA (sgRNA): UGAAAGUUACAACCUCCGCG), which were scored for reduced pigmentation. These injections were compared to injections of 25µM Cas9 nuclease SpCas9-2NLS only, which were subsequently used as experimental controls. For *insm1a* knockdown, 1 nL of solution was injected per embryo that consisted of: 25 µM each of two redundant synthetic guide RNAs targeting *insm1a* (sgRNA1: AGCGACACCUGUUUCGUACC; sgRNA2: CGAACUGCACGGGUGACACC), 25 µM SpCas9-2NLS, and 2.25 ng of tp53 morpholino, all compared to control 25µM SpCas9-2NLS only injections. Synthetic guide RNAs and SpCas9-2NLS were ordered from Synthego (www.synthego.com).

#### Imaging and quantitative analysis

Embryos were prepared and mounted for imaging as previously described^11, 67^. Confocal time-lapse microscopy was performed on a Zeiss LSM 800 microscope with a 40x/1.1 W objective. Data were processed, segmented, and analyzed using Imaris (Bitplane, Inc.) and exported as PNG images or MP4 movies. To assay for changes in Notch signaling activity of Tp1:d2GFP+ cells during the course of the time-lapse, 6 µm spots were used to track cells over time as they migrate apically and their mean fluorescence intensity levels were used to quantify changes in GFP levels over time. For analysis of HCR signal, Imaris’s ‘Spots’ extension was used to manually place 6-8 µm diameter spots on individual cells in the basal region of the OE, using DAPI signal to segment nuclei. Additional spots were placed in other regions of embryos that had no HCR signal to measure the mean background signal intensity for each channel, which was then subtracted from the HCR signal intensity for all spots per channel per embryo. Intensity means were normalized between 0 and 1 across all groups and reported as per cell mRNA levels. Graphs show 1-2 representative datasets chosen from multiple experimental repetitions. For data collected for computational modeling (below), relative *her4.1* and *insm1a* mRNA levels were normalized to *b-actin* mRNA. *b-actin* levels were measured in OE basal cells with two different HCR amplification and fluorophore systems corresponding to those used to perform HCR for *her4.1* and *insm1a*. After background subtraction, data for mean *her4.1* and *insm1a* HCR signal intensities were divided by the corresponding mean *b-actin* signal intensities and normalized to a scale between 0 and 1 across all datasets and time points. For automated signal detection and counting of neurons across control and experimental samples, OEs were selected as regions of interest and 4.5 μm diameter spots were placed with uniform thresholds to detect Ngn1:nRFP+ cells and Ngn1:GFP+ cells. The number of neurons per embryo were normalized to the means of corresponding controls for each experiment. For calculating the fraction of Tp1:d2GFP+ cells in the OE, the ratio of GFP+ cells to that of DAPI+ cells was calculated by automated signal detection and counting for control and experimental samples. To assay for changes in radial migration of Tp1:d2GFP+ cells, the cells with maximum movement during the course of the time-lapse were tracked using the spots feature and the change in their distances from the origin (apical opening of the olfactory cavity) was calculated. To assay for changes in center cells, direct neighbor cells, and indirect neighbor cells, neighborhoods were visually identified for each OE and different cell-types were manually counted for each neighborhood. Calculations to determine statistical significance were performed using Student’s t-test of variables (two-sample t-test assuming unequal variances). Figure legends include n values for experiments.

#### Computational modeling

In our model, the genes *her4.1* and *insm1a* and their products Her4.1 and Insm1a participate in a mutual inhibition network, with RA promoting the production of Insm1a and inhibiting the expression of *her4.1*, while Notch signaling promotes the production of Her4.1 and inhibits the expression of *insm1a* (Fig **4A****, S6A**). The biochemical reactions in the network are listed below:

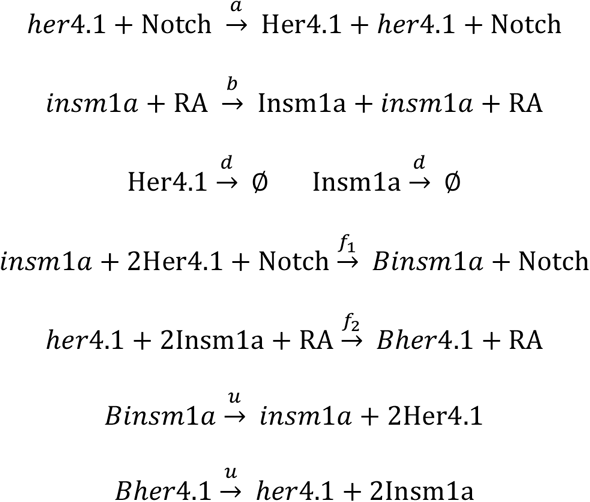

Several stochastic processes are encoded in the underlying reaction network. These reactions describe (1) the synthesis of Her4.1 dependent on Notch signaling and synthesis of Insm1a dependent on RA with reaction rates *a* and *b*, respectively; (2) the degradation of Her4.1 and Insm1a with equal rates *d*; (3) the inhibition of *insm1a* expression by Notch signaling and the inhibition of *her4.1* expression by RA with rates *f*_*1*_ and *f*_*2*_, respectively; (4) the unbinding of Her4.1 to *insm1a* promoter and Insm1a to *her4.1* promoter with equal rates *u*. To approximate experimental results with the reaction network, we measured the ratio between average mRNA levels of *her4.1* and *insm1a*, both normalized to levels of *b-actin* mRNA levels, across time points 24 hpf, 28 hpf, 32 hpf, and 36 hpf and found that ratio to be 0.87. We then set the ratio between rate constants *a* and *b* based on the relative mean mRNA levels of *her4.1* and *insm1a*. To construct a working model, we initially assumed that the strength of Notch signaling’s effect on inhibition of expression of gene *insm1a* is equal to that of the effect of RA signaling on the inhibition of gene expression for *her4.1*. The homodimer binding rates of Insm1 to the promoter region of the gene *her4.1* and Her4.1 to the promoter region of gene *insm1a* are assumed to be approximately 100 times more than the unbinding rate (fast binding). Based on these assumptions, we set the following values for our model parameters: *a* ≈ 0.87*b*; *f*_*1*_ ≈ 0.1*a, f*_*1*_ ≈ *f*_*2*_, *u* ≈ 0.05*f*_*1*_. For the stochastic reaction network (Fig **4A**, **S6A**), we computed the time-evolving probability landscapes of ***p*** = ***p*** (Her4.1, Insm1a) using the previously described ACME Method^30, 31^ as follows:

##### Computing probability landscape using ACME

Assuming that the immediate neighborhood of individual cells is a well-mixed system of reactions with constant volume and temperature, this system has *n* species *X*_*i*_, *i* = 1,2, …, *n*, in which each particle can participate in *m* reactions *R*_*k*_, *k* = 1,2, …, *m*. A microstate of the system at time *t*, ***x***(*t*) is a column vector representing the copy number of species: ***x***(*t*) = (*x*_1_(*t*),*x*_2_(*t*), …, *x*_*n*_(*t*))^*T*^, where the values of copy numbers are non-negative integers. The state space Ω of the system includes all the possible microstates of the system from *t* = 0 to infinity, Ω = {***x***(*t*)|*t* ∈ [0,∞)}.

The reaction *R*_*k*_ of the system takes the form of

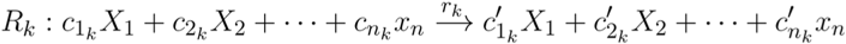

which brings the system from a microstate *x* to a new microstate ***x*** + ***s***_*k*_, where ***s***_*k*_ is the stoichiometry vector and is defined as

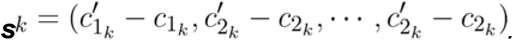

In a well-mixed system, the propensity function of reaction *k, A*_*k*_(***x***) is given by the product of the intrinsic reaction rate constant *r*_*k*_ and possible combinations of the relevant reactants in the current state ***x***.

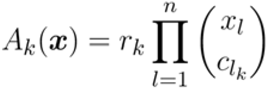

With the above definitions, the discrete Chemical Master Equation (dCME) of a network model of the stochastic chemical kinetic processes consists of a set of linear ordinary differential equations defining the changes in the probability landscape over time at each microstate ***x***. Denoting the probability of the system at a specific microstate ***x*** at time *t* as *p*(***x***,*t*) ∈ R_[0,1]_ and the probability landscape of the system over the whole state space Ω as ***p***(*t*) = {*p*(***x***(*t*))|***x***(*t*) ∈ Ω}, the dCME of the system can then be written as the general form of

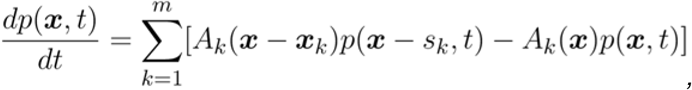

where ***x*** and ***x*** − ***s***_*k*_ ∈ Ω.

The steady state probability landscape is obtained by solving the dCME directly. The exact solution is made possible by using the ACME algorithm as previously described^30-36^. Fig **4B** show the probability landscape of ***p*** = ***p*** (Her4.1, Insm1a) at steady state, which has a multistable configuration and identifies three stable phenotypic states for this network. The first peak where both Her4.1 and Insm1a are inhibited (around Her4.1 = 0 and Insm1a = 0; *her4.1*^OFF^*insm1a*^OFF^) is where the homodimer of Insm1a is bound to the promoter region of gene *her4.1*, and the homodimer of Her4.1 is bound to the promoter region of gene *insm1a*. The second peak is where the expression of *insm1a* is active (around Her4.1 = 0 and Insm1a = 20; *her4.1*^OFF^*insm1a*^ON^). In addition, there is a third peak where *insm1a* is inhibited and *her4.1* is active (around Her4.1 = 15 and Insm1a = 0; *her4.1*^ON^*insm1a*^OFF^). We assume that each gene produces a maximum of 40 proteins in a small region. The copy number of each gene product over the maximum number can be assumed to be the expression level of each gene. The error of the probability modeling is set to 1/295 (296: total number of cells assayed in experiments for mRNA levels from 24 to 36 hpf), as previously described^30, 31^. Probability heatmaps are plotted on a log scale with regions of high probability in green and regions with probability close to zero in white.

